# PHF6 suppresses self-renewal of leukemic stem cells in AML

**DOI:** 10.1101/2024.01.06.573649

**Authors:** Sapana S. Jalnapurkar, Aishwarya Pawar, Subin S. George, Charles Antony, Jason Grana, Sandeep Gurbuxani, Vikram R. Paralkar

## Abstract

Acute myeloid leukemia is characterized by uncontrolled proliferation of self-renewing myeloid progenitors. PHF6 is a chromatin-binding protein mutated in myeloid leukemias, and its loss increases mouse HSC self-renewal without malignant transformation. We report here that *Phf6* knockout increases the aggressiveness of *Hoxa9*-driven AML over serial transplantation, and increases the frequency of leukemia initiating cells. We define the *in vivo* hierarchy of *Hoxa9*-driven AML and identify a population that we term the ‘LIC-e’ (leukemia initiating cells enriched) population. We find that *Phf6* loss has context-specific transcriptional effects, skewing the LIC-e transcriptome to a more stem-like state. We demonstrate that LIC-e accumulation in *Phf6* knockout AML occurs not due to effects on cell cycle or apoptosis, but due to an increase in the fraction of its progeny that retain LIC-e identity. Overall, our work indicates that *Phf6* loss increases AML self-renewal through context-specific effects on leukemia stem cells.

**Statement of Significance:** Leukemia stem cell self-renewal is critical to the pathophysiology of AML. *Phf6* deletion accelerates mouse AML by increasing LSC self-renewal, specifically by increasing the fraction of LSC progeny that retain LSC identity. Our work shows how a repressor of HSC self-renewal is inactivated in AML to drive LSC stemness.

## Introduction

Acute myeloid leukemia (AML) is an aggressive bone marrow malignancy characterized by aberrant proliferation of hematopoietic stem and progenitor cells that are blocked in differentiation. Genomic profiling has revealed that combinatorial somatic mutations in gene classes spanning epigenetic regulators, signaling proteins, transcription factors, and others contribute to leukemogenesis (1). The overall prognosis of AML remains poor, with a need for improved treatments. A deeper understanding of the cellular mechanisms through which AML mutations exert their pathogenic effects is required to advance this goal.

*PHF6* (Plant homeodomain-like finger protein 6) is an X-chromosome gene mutated in a variety of myeloid and lymphoid leukemias. PHF6 protein localizes to the nucleolus and nucleoplasm, and is known to interact with chromatin, but its precise molecular function is poorly understood, with reported roles ranging from cell cycle control (2–4), DNA repair (4,5), and chromatin and transcriptional regulation (6–9). Somatic *PHF6* mutations were first documented in T-cell acute lymphocytic leukemia (T-ALL), where it was found to be mutated in 38% of patients (10). Subsequently, *PHF6* mutations were documented in myeloid malignancies, including 3-6% of AML, myelodysplastic syndrome (MDS), and chronic myelomonocytic leukemia (CMML), as well as in 23% of mixed-phenotype acute leukemia (MPAL) and undifferentiated leukemia (1,11–16). *PHF6* mutations co-occur in MDS/AML with mutations in *RUNX1, ASXL1, and U2AF1* (*1,12,16*), and a majority of *PHF6* mutations are frameshift and nonsense mutations distributed throughout the gene body (16), presumably leading to null alleles and indicating that PHF6 acts as a leukemia suppressor.

Germline *Phf6* deletion in mice leads to perinatal lethality, likely due to non-hematopoietic phenotypes, while mice with hematopoietic *Phf6* deletion are viable and fertile (17,18). Conditional hematopoietic knockouts using multiple *Cre* systems have consistently shown minimal alterations to homeostatic hematopoiesis, but striking increases in HSC self-renewal on transplantation, with the ability to engraft recipient marrow beyond five serial transplants without evidence of exhaustion, malignant transformation, or lineage skewing (17–19). *Phf6* knockout HSCs from aged mice show transcriptional profiles similar to young HSCs, and deletion of *Phf6* from older mice shows a shift towards a younger HSC transcriptome, suggesting that *Phf6* loss may counteract HSC aging (5). Combination of *Phf6* loss with overexpression of activating mutants of *Notch1* (*18*) or *Jak3* (*20*), or overexpression of wild-type *Tlx3* (*17*) has been shown to cause T-ALL acceleration, while transgenic crosses of *Phf6* deletion with *Idh2* mutation produce mixed myeloid-lymphoid leukemias (21). Collectively, the role of PHF6 appears to be the repression of self-renewal, both in normal HSCs as well as in T-ALL (18). However, there are also reports of PHF6 being required for the growth of B-ALL (22), and more recently, for the growth of AML driven by *BCR-ABL*, *AML1-ETO*, or *MLL-AF9* fusions (23). The latter publication reporting the counterintuitive finding that *Phf6* loss reduces AML growth and stemness contradicts the model of PHF6 as a leukemia suppressor; however, the publication’s use of fusion protein drivers that do not co-occur with human *PHF6* mutations may indicate that the chosen AML models recapitulated narrow disease subsets with particular fusion-induced phenotypes, potentially not reflective of broader AML biology. The precise role of *Phf6* loss in AML therefore remains unclear.

In this study, we avoid potential confounding effects of specialized fusion proteins by using *Hoxa9* retroviral transduction (without *Meis1*) as a model of AML (24) that is broadly relevant, given that >70% of AMLs overexpress HOXA9 (25). We examine the role of hematopoietic *Phf6* deletion on AML progression and stemness, and show that *Phf6* loss accelerates AML progression over serial transplantation, accompanied by an accumulation of leukemia initiating cells (LICs). We identify that LICs in the *Hoxa9* transduction model are concentrated within a small Kit+ Ly6C- subpopulation that we term ‘LIC-e’ (LIC-enriched), and we show that, contrary to prior reports, *Phf6* loss has no effect on cell cycle or apoptosis, but instead purely increases the fraction of LIC-e progeny that retain persistent LIC-e identity. We further show that *Phf6* loss leads LIC-e cells to gain a more stem-like transcriptome, along with reduced accessibility of genomic regions bound by the transcription factors AP-1 family, GATA2, and SPI1, among others. Collectively, our data resolves a controversy in the literature by demonstrating that PHF6 suppresses AML stem cell self-renewal (and not cell cycle progression or apoptosis) in a broadly relevant AML model system, and demonstrates how the loss of a specific repressor of HSC self-renewal drives leukemia stemness.

## RESULTS

### PHF6 loss increases leukemia initiating cell frequency in *Hoxa9*-driven AML

To determine the prognostic significance of *PHF6* mutations in human AML, we used publicly available mutational and survival data from the BEAT AML dataset (26). Of 557 AML patients, 16 (2.87%) had *PHF6* mutations. Almost all *PHF6* mutations occurred in patients classified into European Leukemia Net (ELN) intermediate and adverse risk groups. *PHF6* mutations were associated with reduced overall survival in adverse risk patients (median survival 5.56 months in *PHF6* mutated versus 10.16 months in unmutated; *p = 0.016*) (**Fig 1A**) while survival differences did not reach statistical significance in intermediate-risk patients, potentially due to relatively small case numbers (median survival 9.21 months in *PHF6* mutated versus 12.73 months in unmutated; *p = 0.26*) (**Fig S1A**). The dataset was not powered for further multivariable or co-mutation analyses, but overall indicated that *PHF6* mutation is associated with worse survival.

**Figure 1.**
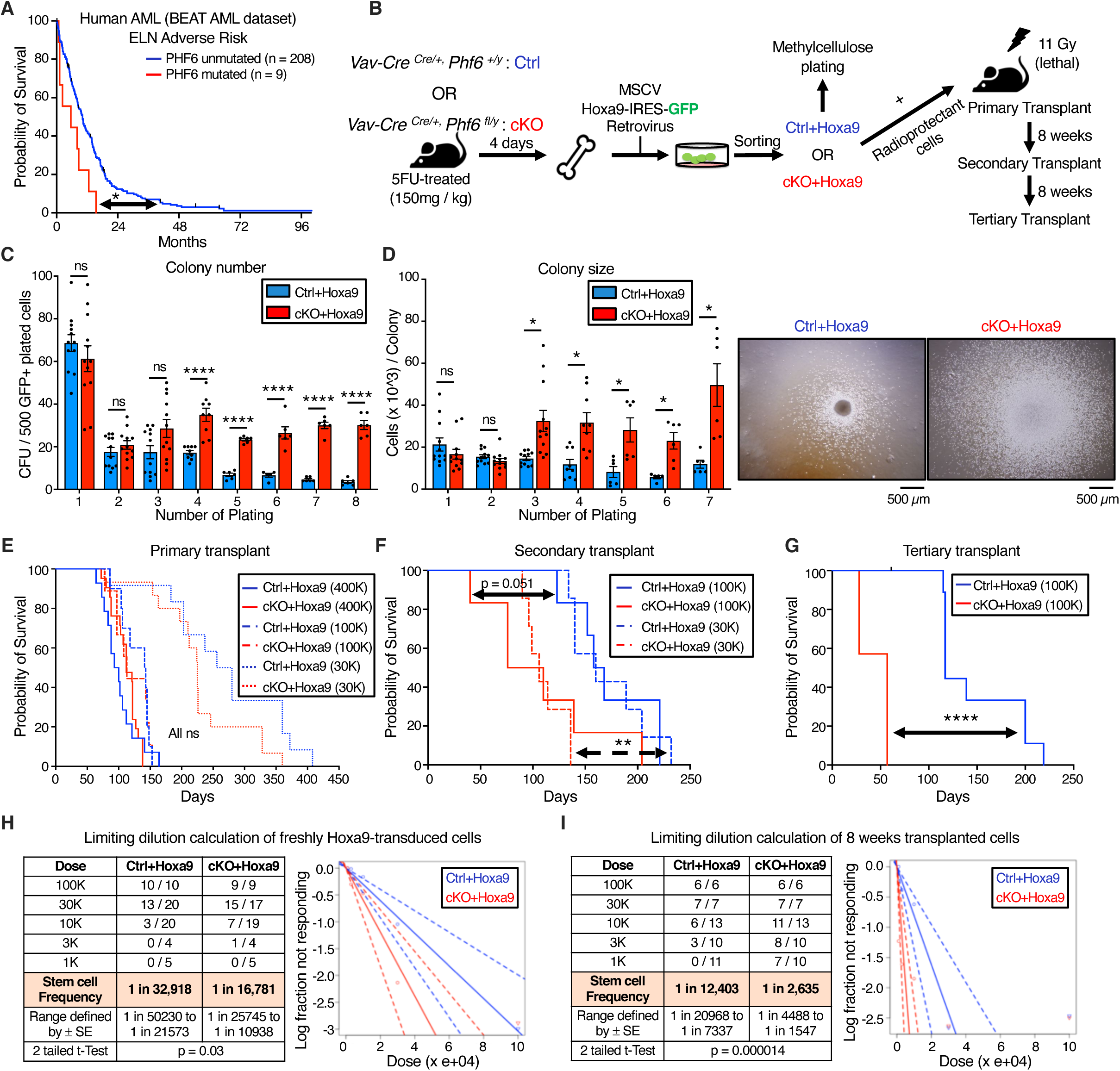
*Phf6* loss increases leukemia initiating cell frequency in *Hoxa9-*driven AML. **A,** Kaplan-Meier survival curve for *PHF6* mutated and unmutated adverse risk adult AML patients (ELN classification) from the BEAT AML dataset. **B,** Experimental design for *Hoxa9* retroviral transduction of *Ctrl* and *cKO* marrow, followed by colony forming unit assay (CFU), AML induction in mice, and serial transplantation of transformed leukemic cells. **C,** Bar graph showing number of colony forming units (CFUs) obtained from 8 rounds of serial methylcellulose replating of 500 cells/plate of *Ctrl+Hoxa9* and *cKO+Hoxa9* transformed mouse bone marrow. (n = 6-12 biological replicates) **D,** *Left* Bar graph showing average number of cells per colony (colony size) obtained after 7 rounds of serial methylcellulose replating of 500 cells/plate of *Ctrl+Hoxa9* and *cKO+Hoxa9* transformed mouse bone marrow. (n = 6-12 biological replicates) *Right* Representative photographs of colonies at the 3^rd^ plating. Scale bar represents 500 μm. **E,** Kaplan-Meier survival curves of *Ctrl+Hoxa9* and *cKO+Hoxa9* primary transplant recipients receiving 400K, 100K or 30K GFP+ cells. (n = 7-21 mice per cohort) **F,** Kaplan-Meier survival curve of *Ctrl+Hoxa9* and *cKO+Hoxa9* secondary transplant recipients, receiving 100K or 30K GFP+ cells harvested from bone marrow of primary recipients 8 weeks after transplantation. (n = 7-10 mice per cohort) **G,** Kaplan-Meier survival curve of *Ctrl+Hoxa9* and *cKO+Hoxa9* tertiary transplant recipients, receiving 100K GFP+ cells harvested from bone marrow of secondary recipients 8 weeks after transplantation. (n = 7-11 mice per cohort) **H** and **I,** Limiting dilution analysis for LIC calculation for (**H**) freshly *Hoxa9*-transduced cells, (**I**) 8 weeks primary transplanted leukemic cells. All bar graphs show mean ± SEM, and statistical significance was calculated using the Student t-test. For all survival curves, statistical significance was calculated using the Log-rank (Mantel-Cox) test. *\*p < .05, **p < .01, ***p < .001; ****p < .0001, ns = not significant*.

To assess the role of PHF6 loss in mouse AML, we generated conditional hematopoietic *Phf6* knockout *Vav-Cre^Cre/+^Phf6^fl/y^* (*cKO*) mice and compared them to their *Vav-Cre^Cre/+^Phf6^+/y^* (*Ctrl*) littermates. *cKO* marrow was confirmed to show loss of *Phf6* mRNA (**Fig S1B**). Published studies of hematopoietic *Phf6* loss have reported no evidence of leukemic transformation either in homeostatic mice or over serial bone marrow transplantations (17–19,27). Consistent with these reports, we found no blood count abnormalities in *cKO* mice compared to *Ctrl* mice up to 9 months of age (**Fig S1C-I**), further indicating that *Phf6* loss alone is likely insufficient to initiate overt leukemia.

We next induced AML using the *Hoxa9* retroviral transduction model (*Hoxa9* alone, without *Meis1*). We picked this model due to its ability to produce AML with a relatively longer latency (lethality in ∼3-6 months) (24), allowing us to test a potential role for *Phf6* loss in accelerating AML kinetics. Given that 70% of human AMLs show high *HOXA9* levels (25), the *Hoxa9* overexpression mouse model is broadly relevant to human leukemia biology, and avoids caveats associated with specific fusion proteins. We transduced whole bone marrow from 5-FU-treated *Ctrl* and *cKO* mice with *MSCV Hoxa9*-*IRES-GFP* retrovirus, and first investigated the effect of *Phf6* loss on the ability of *Hoxa9*-transformed cells to form colonies in methylcellulose (**Fig 1B**). We observed that *Ctrl+Hoxa9* cells were nearly exhausted after 4 platings, whereas *cKO+Hoxa9* cells demonstrated persistent colony-forming ability up to 8 platings (**Fig 1C**), with larger colonies (**Fig 1D**). Thus, *Phf6* loss gives a replating advantage to *Hoxa9*-transformed marrow and delays its *in vitro* exhaustion.

To test the role of *Phf6* loss in the development of AML *in vivo*, we transplanted *Hoxa9*-transduced marrow into lethally irradiated syngeneic recipients (**Fig 1B)**. While previous publications have reported AML development with *Hoxa9*-only transduction (24), detailed histological characterization of this particular model is lacking in the literature. We therefore first confirmed that *Ctrl+Hoxa9* marrow produced lethality in recipient mice in ∼3-5 months after transplantation (**Fig 1E)** due to AML characterized by accumulation of >20% blasts in the marrow (**Fig S1J**), peripheral leukocytosis with circulating blasts (**Fig S1K**), splenomegaly with effacement of splenic architecture (**Fig S1L**), and infiltration of leukemic cells in the liver (**Fig S1M**). Survival was similar between *Ctrl+Hoxa9* and *cKO+Hox*a9 groups in primary recipients transplanted with multiple doses (400K, 100K, or 30K cells) (**Fig 1E)**, with similar degrees of leukemic infiltrate (**Fig S2A-F**) and splenomegaly at morbidity **(Fig S2G)**. However, serial transplantation (secondary and tertiary) of bone marrow showed accelerated lethality in *cKO+Hoxa9* compared to *Ctrl+Hoxa9* (**Fig 1F-G, S2H**), with median survivals of 106 days versus 160 days in secondary recipients (*p = 0.001*) and 57 days versus 117 days in tertiary (*p < 0.0001*). Thus, loss of *Phf6* accelerates *Hoxa9-*driven mouse AML on serial transplantation.

We next sought to determine the effect of *Phf6* loss on the frequency of leukemia initiating cells (LIC), the sub-population of transformed marrow capable of initiating leukemia. We performed limiting dilution transplantation assays (LD) on freshly transduced marrow (GFP+ cells sorted 2 days after initial retroviral transduction) as well as on marrow from transplant recipients (GFP+ bone marrow cells sorted from recipients 8 weeks after primary transplantation). We picked the 8-week time point based on the initiation of lethality in this model at ∼12 weeks (**Fig 1E**), allowing us to harvest cells from all transplanted mice at a standard time point prior to onset of lethality. LD of freshly transduced marrow showed that, at baseline, *cKO+Hoxa9* cells had a 2-fold higher frequency of cells capable of leukemic transformation (1 in 16,781 versus 1 in 32,918, *p = 0.03*) (**Fig 1H**). LD on marrow extracted 8 weeks post-transplantation showed an approximately 5-fold greater frequency of leukemia initiating cells (LICs) in *cKO+Hoxa9* marrow, with LIC frequency of 1 in 2,635 versus 1 in 12,403 for *Ctrl+Hoxa9* marrow (*p = 0.000014*) (**Fig 1I**). Thus, *Phf6* loss increases LIC frequency in *Hoxa9*-driven AML, with a progressive increase in LIC frequency during *in vivo* AML evolution.

### *Phf6* loss increases leukemic disease burden

To characterize the effect of *Phf6* loss further, we analyzed peripheral blood, splenic architecture, and bone marrow leukemic cell burden of primary recipient mice at 8 weeks after transplantation. Mice transplanted with *cKO+Hoxa9* cells showed a higher frequency of GFP+ cells in peripheral blood at 8 weeks than mice receiving *Ctrl+Hoxa9* cells (**Fig 2A**). The *cKO+Hoxa9* group also had greater leukocytosis (**Fig 2B**) and more severe thrombocytopenia (**Fig 2C**). Mice in both groups displayed comparable levels of anemia (**Fig S3A-B**). The *cKO+Hoxa9* group had increased spleen size and weight compared to the *Ctrl+Hoxa9* group (**Fig 2D-E),** and histopathological analysis showed greater effacement of splenic architecture (**Fig 2F**). Splenic infiltration was quantified using a previously described leukemia infiltration score (28), and was found to be greater in *cKO+Hoxa9* mice compared to *Ctrl+Hoxa9* (**Fig 2G**). Giemsa stained cytospin preparations showed higher blast percentages in *cKO+Hoxa9* at the 8-week timepoint (**Fig 2H-I**), and flow cytometry showed higher absolute and percent GFP+ cells (**Fig 2J, S3C**). As expected, virtually all GFP+ cells were myeloid in both groups (**Fig 2K, S3D**). Collectively, though mice from both groups succumbed at similar times after primary transplant (**Fig 1F**), analyses at matched time points before the onset of mortality revealed greater disease burden in *cKO+Hoxa9* mice compared to *Ctrl+Hoxa9*.

**Figure 2.**
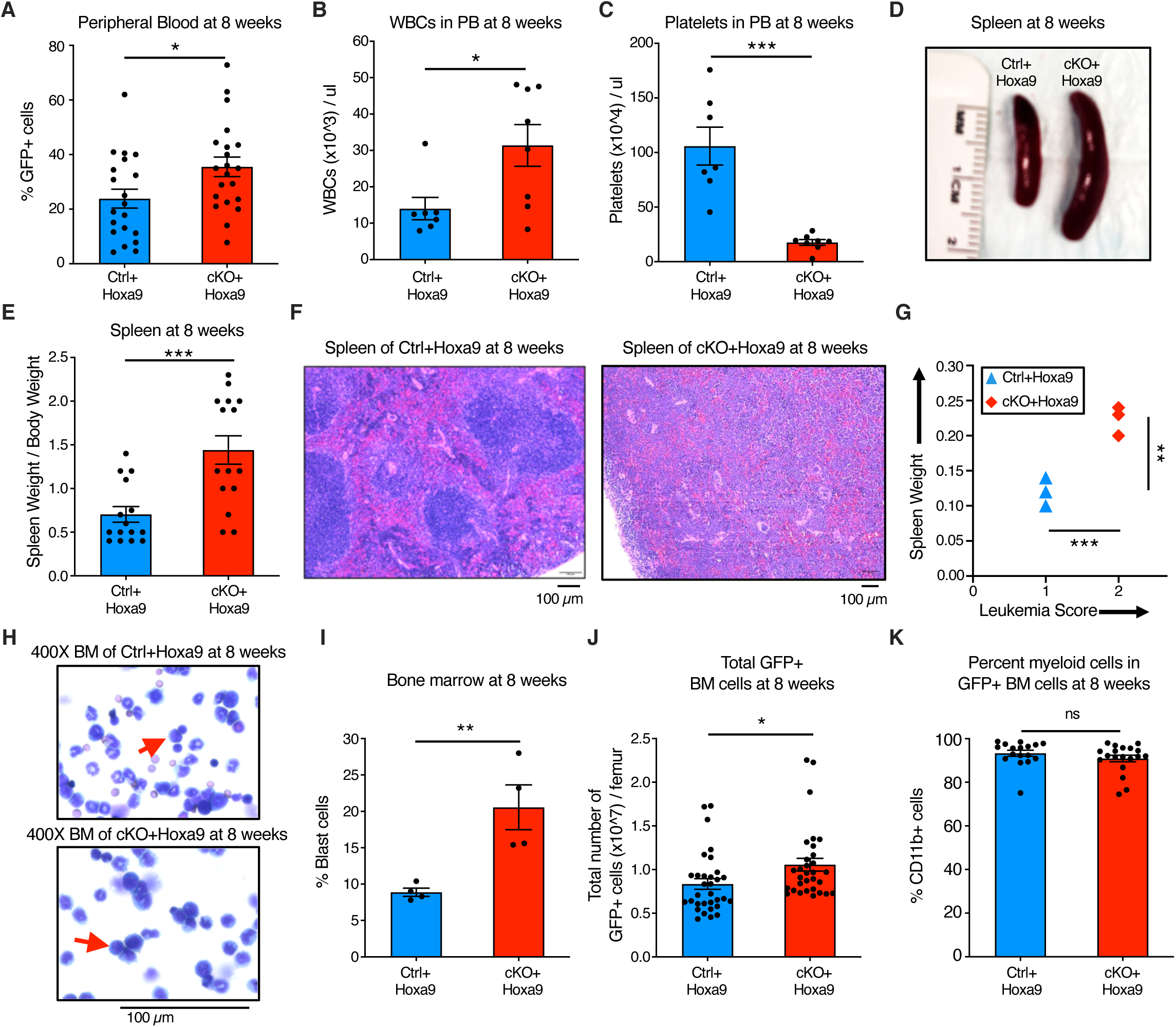
*Phf6* loss increases leukemic disease burden. **A, B,** and **C,** Bar graphs showing peripheral blood analysis at 8 weeks after transplantation of *Ctrl+Hoxa9* and *cKO+Hoxa9* cells. (**A**) Percentage of GFP+ cells in peripheral blood. Counts of (**B**) WBCs and (**C**) platelets in peripheral blood. Normal range for WBCs: 2000-10,000/µl. Normal range for platelets: 900-1,600 x 10^3^/µl (56). **D,** Representative photograph of spleens at 8 weeks after transplantation. Ruler depicts length in centimeters. **E,** Bar graph showing spleen weight (expressed as percentage of total body weight) of primary recipients at 8 weeks after transplantation of *Ctrl+Hoxa9* and *cKO+Hoxa9* cells. **F,** Representative image of H&E staining of spleen from *Ctrl+Hoxa9* and *cKO+Hoxa9* at 8 weeks after transplantation. Scale bar is 100µm at 10X. **G,** Spleen weight (y-axis) and leukemia score (x-axis) from *Ctrl+Hoxa9* and *cKO+Hoxa9* primary recipients at 8 weeks. X-axis represents a previously published leukemia infiltration score (28) calculated based on splenic architecture. Intact white and red pulp was scored as 0, extramedullary hematopoiesis evident by aberrant cells in disturbed white pulp was scored as 1, while infiltration with leukemic blasts with high mitotic activity was scored as 2. **H,** Representative image of Wright-Giemsa staining of cytospin of *Ctrl+Hoxa9* and *cKO+Hoxa9* bone marrow cells at 8 weeks after transplantation. Scale bar is 100µM at 400X. Red arrows indicate representative blast cells. **I,** Bar graph of percentage blast cells of total nucleated cells in bone marrow at 8 weeks after transplant. **J** and **K,** Bar graphs showing (**J**) absolute number of GFP+ cells in marrow and (**K**) percentage of CD11b+ myeloid cells among all GFP+ cells at 8 weeks after transplantation. All bar graphs show mean ± SEM, and statistical significance was calculated using the Student t-test. For all survival curves, statistical significance was calculated using the Log-rank (Mantel-Cox) test. *\*p < .05, **p < .01, ***p < .001; ****p < .0001, ns = not significant*.

### *Phf6* loss increases the frequency of self-renewing, transplantable LICs

To characterize the immunophenotype of AML subpopulations (including LICs), we further analyzed the bone marrow of *Ctrl+Hoxa9* recipients at 8 weeks after transplantation. As expected, GFP+ cells did not express B or T cell markers (**Fig S4A**). Immature AML cells are known to have high c-Kit expression (29), and leukemic stem cells (LSCs) in the MLL-AF9 retroviral mouse model aberrantly express mature myeloid lineage antigens such as Ly6C and CD11b (30). To our knowledge, corresponding AML subpopulations have not been defined in *Hoxa9*-only-driven AML. To identify the subpopulation containing LICs, and to characterize the differentiation hierarchy of *Hoxa9*-driven AML, we tested multiple gating strategies with a panel of markers, eventually settling on a simplified strategy using c-Kit and Ly6C expression to divide GFP+ bone marrow cells into three populations: i) cKit+ Ly6C-, ii) c-Kit+ Ly6C+, and iii) c-Kit- Ly6C+ (**Fig 3A, S4B**). We established the hierarchy of these populations based on differentiation trajectory, immunophenotype, colony-forming ability, and *in vivo* repopulation capacity. The population at the top of the hierarchy was the cKit+ Ly6C- population, an immature population with expression of cKit, CD34, and dim CD11b, with no expression of Ly6C, Ly6G, or Sca-1, and mixed expression of CD16/32 (**Fig S4C**). On sorting and culture, this population was capable of giving rise to more differentiated Ly6C+ cells within 2 days (**Fig S4D**), could produce colonies on methylcellulose plating (**Fig 3B**), and could engraft into recipient mice (**Fig 3C**). Based on this subpopulation’s ability to engraft, but cognizant that not all cells within it are LICs, we termed it the ‘LIC enriched’ (LIC-e) population (**Fig 3D**). The second population was the c-Kit+ Ly6C+ population, also expressing CD11b, CD34, and CD16/32, but not Sca-1 (**Fig. S4C**). On culture, this population could only give rise to Ly6C+ cells, but not to any Ly6C- cells (**Fig. S4D**), indicating that it is irreversibly committed to differentiation. This population could produce a small number of colonies in methylcellulose (**Fig 3B**), but could not engraft mice (**Fig 3C**). We termed it the ‘committed’ leukemic cell population (**Fig 3D**). The third population was the c-Kit- Ly6C+ population, which expressed Ly6C, CD11b, CD34, and CD16/32, but not Sca-1, and had mixed expression of Ly6G (**Fig S4C**). It could not produce any colonies (**Fig 3B**) nor engraft (**Fig 3C**). In vitro, this population only gave rise to Ly6C+ cells (**Fig. S4D**). We termed it the ‘differentiated’ leukemic cell population (**Fig 3D**). We thus established the hierarchical organization of *Hoxa9*-driven AML, and used it to evaluate the effects of *Phf6* loss.

**Figure 3.**
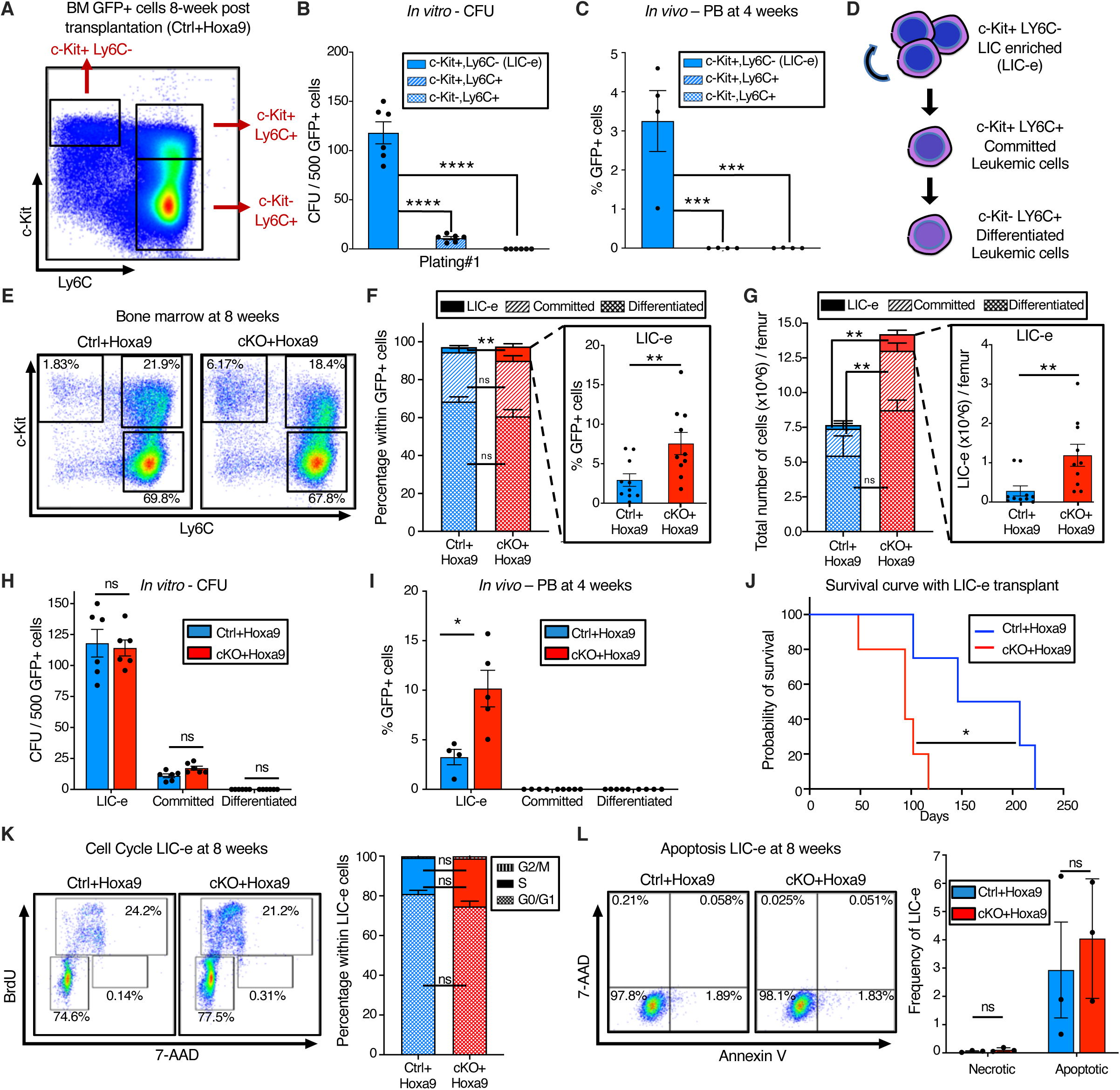
*Phf6* loss increases the frequency of self-renewing, transplantable LICs. **A,** Representative flow cytometry plot of bone marrow GFP+ cells at 8 weeks after transplantation with *Ctrl+Hoxa9* cells, compartmentalized into three subpopulations i) LIC-e, ii) Committed, and iii) Differentiated leukemic cells using c-Kit and Ly6C expression. Note: The same flow cytometry plot has been shown in Fig S4 with detailed immunophenotypic markers. **B,** Bar graph showing number of colony forming units (CFUs) obtained on methylcellulose plating of 500 cells of sorted subpopulations of *Ctrl+Hoxa9* transplanted marrow. (n = 6 biological replicates) **C,** Bar graph showing frequencies of GFP+ cells in peripheral blood of secondary recipient mice at 4 weeks after transplantation of sorted subpopulations from Ctrl+Hoxa9 primary recipient marrow. (n = 4-5 biological replicates) **D,** Schematic of hierarchical organization of leukemic cells (LIC-e, committed, and differentiated leukemic cells) within AML produced by retroviral *Hoxa9* transduction. **E,** Representative flow cytometry plots depicting subpopulations of *Ctrl+Hoxa9* and *cKO+Hoxa9* leukemia marrow at 8 weeks after primary transplant. **F and G,** Stacked bar graphs showing (**F**) frequencies, and (**G**) absolute number per femur of LIC-e, committed, and differentiated leukemic populations from *Ctrl+Hoxa9* and *cKO+Hoxa9* marrow at 8 weeks after transplantation. Insets show frequencies and absolute numbers of the LIC-e subpopulation. (n = 10-11) **H,** Bar graph showing number of CFUs obtained on methylcellulose plating of 500 cells of sorted LIC-e, committed, and differentiated leukemic populations from *Ctrl+Hoxa9* and *cKO+Hoxa9* primary recipient bone marrow at 8 weeks after transplantation. (n = 6 biological replicates) **I,** Bar graph showing frequencies of GFP+ cells in the peripheral blood of secondary recipient mice at 4 weeks after transplantation with sorted LIC-e, committed, and differentiated leukemic cell subpopulations from *Ctrl+Hoxa9* and *cKO+Hoxa9* primary recipient bone marrow at 8 weeks after transplantation. (n = 5 biological replicates). Note: The *Ctrl+Hoxa9* samples in Figs 3H and 3I are the same as those depicted in 3B and 3C - the current figures are comparing *Ctrl* and *cKO*. **J,** Kaplan-Meier curve of secondary transplant recipients receiving 50K sorted LIC-e cells from *Ctrl+Hoxa9* and *cKO+Hoxa9* primary mouse bone marrow at 8 weeks after transplantation. (n = 4-5 biological replicates) **K,** *Left* Representative flow cytometry plots for cell cycle analysis of LIC-e cells, with BrdU (marking cells in S phase) and 7-AAD (marking DNA) 2 hours after BrdU injection into live mice at 8 weeks after transplantation. *Right* Stacked bar graph indicates frequencies of *in vivo* LIC-e cells in G0/G1, S, and G2/M phases. (n = 13 biological replicates) **L,** *Left* Representative flow cytometry plots of apoptotic (Annexin V+, 7AAD-), and necrotic (7AAD+) LIC-e cells. *Right* Bar graph shows frequencies of apoptotic and necrotic LIC-e cells from *Ctrl+Hoxa9* and *cKO+Hoxa9* primary recipient bone marrow at 8 weeks after transplantation. All bar graphs show mean ± SEM, and statistical significance was calculated using the Student t-test. For all survival curves, statistical significance was calculated using the Log-rank (Mantel-Cox) test. *\*p < .05, **p < .01, ***p < .001; ****p < .0001, ns = non significant*.

The LIC-e population, though comprising a small minority of GFP+ cells, was expanded in 8-week *cKO+Hoxa9* bone marrow compared to *Ctrl+Hoxa9*, while the relative fractions of committed and differentiated leukemic cells were similar (**Fig 3E-F**). The difference in the LIC-e population was even more pronounced when absolute cell numbers were considered, showing a 5-fold increase in *cKO+Hoxa9* (**Fig. 3G**). To determine functional differences between *Ctrl+Hoxa9* and *cKO+Hoxa9* subpopulations, we sorted equal numbers of cells from each subpopulation at 8 weeks after transplant, and performed methylcellulose culture and secondary transplantation into irradiated recipients. Committed and differentiated leukemic cells of either group formed few to no colonies, while LIC-e cells showed comparable colony-forming ability (**Fig 3H**). *cKO+Hoxa9* LIC-e cells showed greater engraftment in secondary recipients at 4 weeks (**Fig 3I)** and led to more rapid lethality than *Ctrl+Hoxa9* LIC-e cells (94 days versus 177 days, *p = 0.02*) (**Fig 3J**). We did not observe any difference in cell cycle distribution or apoptosis of LIC-e cells (**Fig 3K-L**). Thus, *Phf6* loss led to an expanded and more transplantable LIC-enriched AML subpopulation whose enhanced leukemic potential was not explained by differences in cell cycle or apoptosis.

### *Phf6* loss promotes a stemness gene network

We next sought to determine the transcriptional consequences of *Phf6* loss by performing bulk RNA-Seq on LIC-e and committed leukemic cells from bone marrow of transplanted recipients at 8 weeks. Committed leukemic cells showed virtually no change in gene expression as a consequence of *Phf6* loss (**Fig S5A-B, Table S2**), while the LIC-e population showed a focused number of differentially expressed genes, with 91 downregulated and 65 upregulated genes in *cKO+Hoxa9* compared to *Ctrl+Hoxa9* (**Fig 4A-B, Table S2**). Genes downregulated in *cKO+Hoxa9* LIC-e cells included known regulators of self-renewal and differentiation like *Runx1 and Cd34* (*31,32*) and leukocyte maturation genes like *Selplg* (*33*) (**Fig 4B-C**). Genes upregulated in *cKO+Hoxa9* LIC-e *cells* included the thrombopoietin receptor *Mpl*, known to contribute to AML pathogenesis (34,35), *F2r,* a G-protein coupled receptor that promotes stemness in MLL-AF9 AML (36)*, Hemgn,* which promotes self-renewal and myeloid progenitor expansion (37), and *Samd14*, which promotes c-Kit signaling and hematopoietic progenitor expansion (38) (**Fig 4B**). Gene set enrichment analysis (GSEA) (39) showed that the *cKO+Hoxa9* LIC-e transcriptome showed positive enrichment for gene sets related to high LSC potential in MLL-AF9-driven mouse AML (40) and leukemic GMPs (L-GMPs) (41), and negative enrichment for gene sets related to myeloid differentiation (42), low LSC potential (40), mature neutrophils, and mature monocytes (43) (**Fig 4D, S5C)**. Thus, *Phf6* loss has context-specific transcriptional effects, skewing the transcriptome of LIC-e cells to a more stem-like and less differentiated state, but showing no effect in more differentiated leukemic cells.

**Figure 4.**
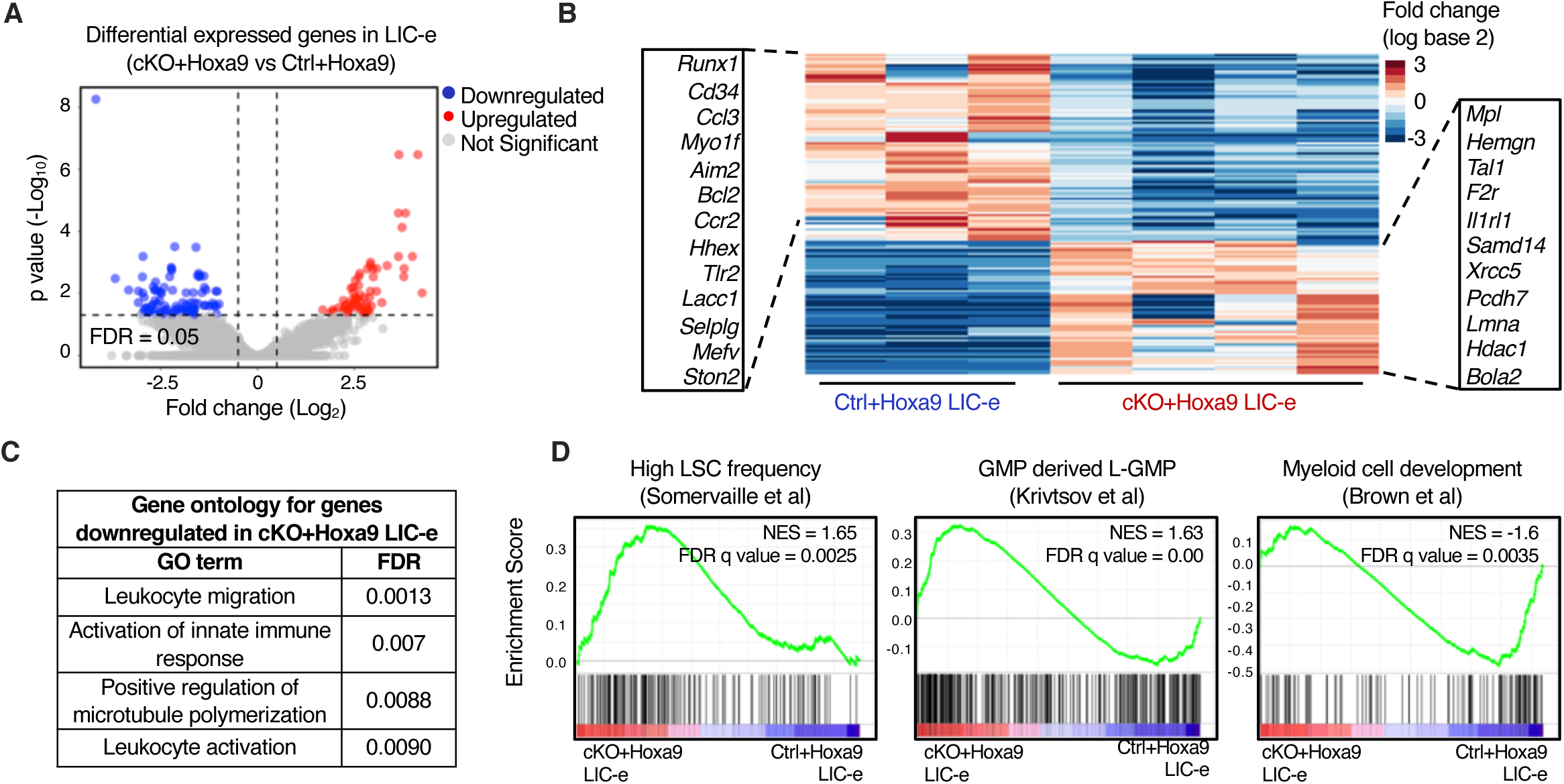
*Phf6* loss promotes a stemness gene network. **A,** Volcano plot showing differentially expressed genes in LIC-e cells from *cKO+Hoxa9* compared to *Ctrl+Hoxa9* bone marrow at 8 weeks after transplantation. (n = 3-4 biological replicates) **B,** Heatmap of differential expression between *Ctrl+Hoxa9* and *cKO+Hoxa9* LIC-e cells. Insets show selected downregulated (left) and upregulated (right) genes in *cKO+Hoxa9* LIC-e compared with *Ctrl+Hoxa9* LIC-e. **C,** Top Gene Ontology terms enriched in genes downregulated in *cKO+Hoxa9* LIC-e compared with *Ctrl+Hoxa9* LIC-e. **D,** Gene set enrichment analysis (GSEA) plots of the *cKO+Hoxa9* LIC-e transcriptome compared to *Ctrl+Hoxa9*. Plots show positive enrichment of gene sets related to high LSC frequency (left) and leukemic GMPs (middle), and negative enrichment of a gene set related to myeloid development (right). Normalized Enrichment scores (NES) and FDR q values are shown.

### *Phf6* loss prevents exhaustion of LIC-e cells by maintaining their self-renewal potential

To determine the kinetics of *Phf6* loss on the behavior of LIC-e cells, we cultured *Hoxa9*-transduced bone marrow in cytokine-supplemented media. The growth rate of bulk culture was similar for *Ctrl+Hoxa9* and *cKO+Hoxa9* marrow (**Fig S6A**). When sorted LIC-e cells were cultured, most cells lost LIC-e identity within days, and the culture was largely replaced by committed and differentiated cells by a week (**Fig 5A**). However, though both groups produced similar fractions of committed (c-Kit+ Ly6C+) and differentiated cells (c-Kit- Ly6C+), the *Ctrl+Hoxa9* culture almost completely depleted its LIC-e population (<1%), while the *cKO+Hoxa9* culture maintained this population, plateauing at 5-6% of the total culture after 5 days (**Fig 5A**). Thus, even though *Phf6* loss does not impair the ability of cultured cells to proliferate and produce differentiated cells, it prevents exhaustion of the LIC-e population, recapitulating the *in vivo* LIC-e accumulation phenotype shown earlier (**Fig 3E-G**).

**Figure 5:**
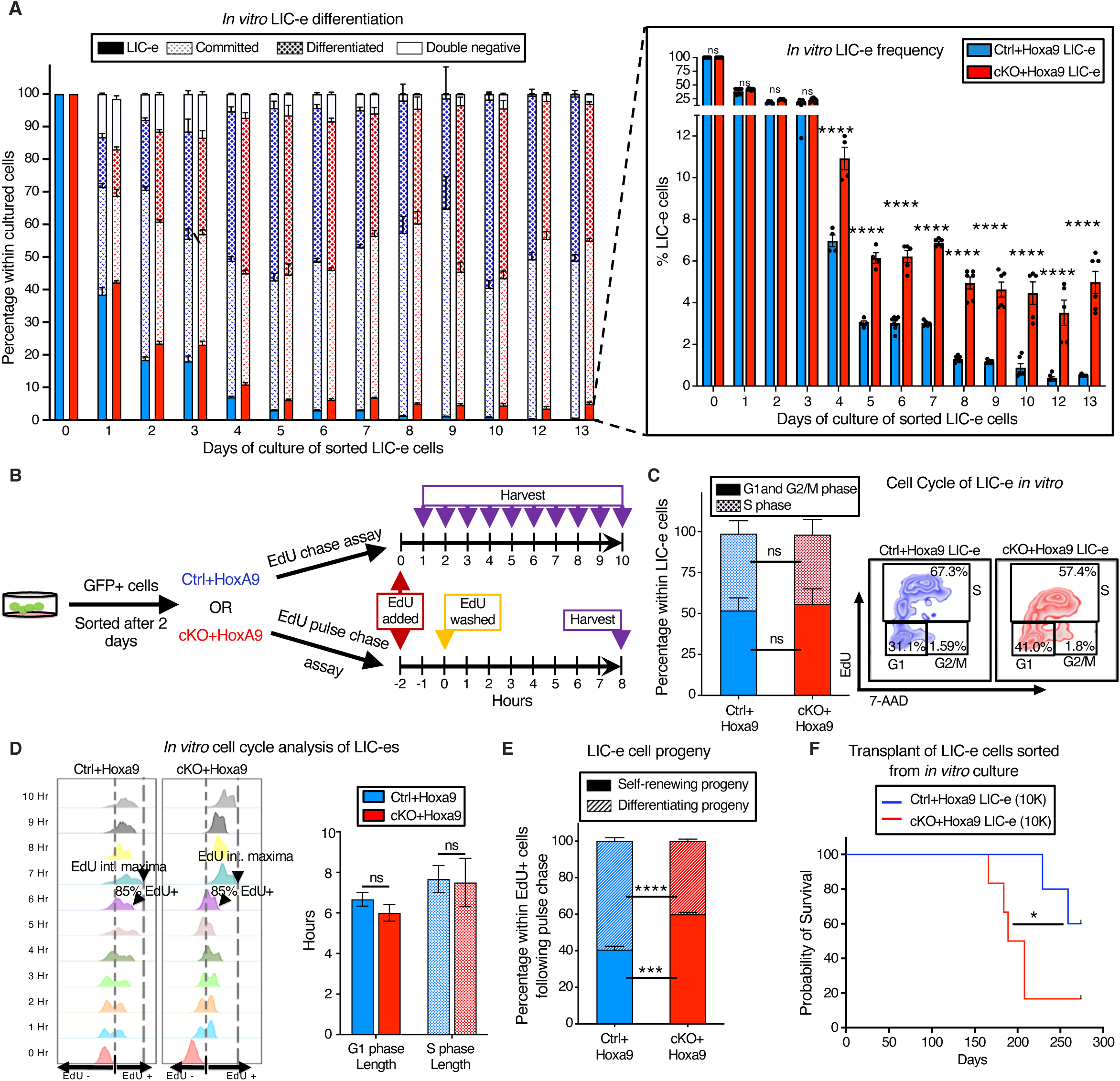
*Phf6* loss prevents exhaustion of LIC-e cells by maintaining their self-renewal potential. **A,** Bar graph showing frequencies of subpopulations resulting from *in vitro* culture of LIC-e cells sorted 4 days after *Hoxa9* transduction of *Ctrl* and *cKO* bone marrow. Inset bar graph depicts only LIC-e frequencies in the same culture. (n = 13 biological replicates) **B,** Experimental design for study of *in vitro* cell cycle analysis (top) and self-renewal (bottom) of LIC-e cells using Edu chase and pulse-chase assay respectively. **C,** *Left,* Bar graph showing frequencies of G0/G1, S, and G2/M phases in *Ctrl+Hoxa9* and *cKO+Hoxa9* LIC-e cells in culture 2 hours after addition of EdU. (n = 4-5 biological replicates) *Right,* Representative flow cytometry plots of same, with EdU marking cells in S phase and 7-AAD staining DNA. **D,** *Left,* Representative flow cytometry plots showing kinetics of uptake of EdU by *Ctrl+Hoxa9* and *cKO+Hoxa9* LIC-e cells over 10 hours of culture, performed for calculation of cell cycle length. Time to whole population (>85%) EdU uptake represents G1 phase length, and time to EdU intensity maxima represents S phase length. *Right,* Bar graph represents the length of cell cycle phases of LIC-e cells in culture. (n = 4-5 biological replicates) **E,** Stacked bar graph showing the percentage of self-renewing and differentiating progeny produced by *Ctrl+Hoxa9* and *cKO+Hoxa9* LIC-e cells in culture. (n = 4-5 biological replicates) **F,** Kaplan-Meier curve of primary transplant recipients receiving 10K sorted *Ctrl+Hoxa9* and *cKO+Hoxa9* LIC-e cells. (n = 5-6 biological replicates) All bar graphs show mean ± SEM, and statistical significance was calculated using the Student t-test. For all survival curves, statistical significance was calculated using the Log-rank (Mantel-Cox) test. *\*p < .05, **p < .01, ***p < .001; ****p < .0001, ns = non significant*.

We had already determined that *in vivo* LIC-e cells showed no difference in cell cycle distribution (**Fig 3K**), but to confirm that LIC-e accumulation wasn’t due to subtle cycling differences, we performed a 10-hour cell cycle analysis in culture by adding EdU to sorted *Ctrl+Hoxa9* and *cKO+Hoxa9* LIC-e cells and harvesting them at serial time points (**Fig 5B**). We observed that the distribution of *Ctrl+Hoxa9* or *cKO+Hoxa9* LIC-e cells in G1 and S phases showed no difference at the start of culture, and showed no difference in the kinetics or magnitude of EdU incorporation (**Fig 5C**). Based on previously published rationale (44) that cells that had just begun G1 would be the last to enter S phase and incorporate EdU, and that the maximum EdU fluorescence intensity reached by the population as a whole would represent cells that had incorporated EdU into their entire genomes, we defined the time required for 85% of each sample to become EdU+ as the ‘G1 phase length’, and the time required for each sample to reach a maximal EdU fluorescent intensity as the ‘S phase length’. We did not observe any difference in G1 and S phase lengths between *Ctrl+Hoxa9* and *cKO+Hoxa9* LIC-e cells (**Fig 5D**), further demonstrating that the phenotype of LIC-e accumulation (**Fig 5A**) is not explained by increased cycling of that subpopulation. We therefore hypothesized that *cKO+Hoxa9* LIC-e cells, while capable of producing differentiated progeny, may have a greater tendency to produce progeny with persistent LIC-e identity. To test this hypothesis, we performed an EdU pulse-chase experiment by incubating sorted LIC-e cells with EdU for 2 hours (pulse), followed by washing off the EdU and further culturing cells for 8 more hours (chase, for the length of S phase) (**Fig 5B**). The goal was to allow all cells that were in S phase during the initial EdU pulse to complete mitosis, so that all EdU+ cells at the end of the 8 hour EdU-free chase would be daughter cells/progeny of the original EdU-uptaking cells. We determined the percentage of self-renewing progeny by calculating the percentage of total progeny (total EdU+ cells) that had LIC-e markers (Ly6C- EdU+), while the rest (Ly6C+ EdU+) were differentiating progeny. We observed that while 40.6% of the progeny of *Ctrl+Hoxa9* LIC-e cells were also LIC-e cells, this fraction was increased to 60.0% in *cKO+Hoxa9* (**Fig 5E**). To confirm that *in vitro* LIC-e cells are functionally equivalent to their *in vivo* counterparts, we performed primary transplantation with 10K LIC-e cells sorted after 2 days of culture. Recipients of *cKO+Hoxa9* LIC-e cells succumbed faster than *Ctrl+Hoxa9* LIC-e (*p = 0.047*) (**Fig 5F**). Collectively, *Phf6* loss prevents the exhaustion of LIC-e cells by increasing the fraction of their progeny that retain persistent LIC-e identity.

### Effects of Phf6 loss on chromatin accessibility in LIC-e cells

To profile the effects of *Phf6* loss on the accessibility landscape of LIC-e cells, we performed ATAC-Seq on sorted LIC-e cells from freshly transduced *cKO+Hoxa9* and *Ctrl+Hoxa9* marrow. We observed that *cKO+Hoxa9* LIC-e cells showed a surprising global reduction in chromatin accessibility compared to *Ctrl+Hoxa9*, with only a few regions showing increased accessibility (**Fig 6A-B**). Regions that lost accessibility in *cKO+Hoxa9* LIC-e cells showed enrichment for AP-1, HOX, SPI, and GATA family motifs, among others (**Fig 6C**). Public ChIP-Seq tracks from leukemic or myeloid cells showed co-occupancy of these factors at ATAC peaks with reduced accessibility (**Fig 6D, Table S3**). Promoters of multiple genes like *Runx1, Selplg,* and *Aim2,* which are downregulated in *cKO+Hoxa9* LIC-e cells (**Fig 4B**), showed occupancy by these factors and showed reduced chromatin accessibility in *cKO+Hoxa9* LIC-e cells (**Fig S6B**). Conversely, the small number of regions that gained accessibility in *cKO+Hoxa9* LIC-e cells showed enrichment for NF-kB, and IRF family motifs (**Fig 6E**). Public ChIP-Seq tracks showed co-occupancy of NF-kB (RELA, RELB) and IRF8, IRF4 at regions of increased accessibility (**Fig 6F, Table S3**). Overall, *Phf6* loss, likely via a combination of direct and indirect effects, led to altered accessibility at sites bound by key hematopoietic transcription factors.

**Figure 6:**
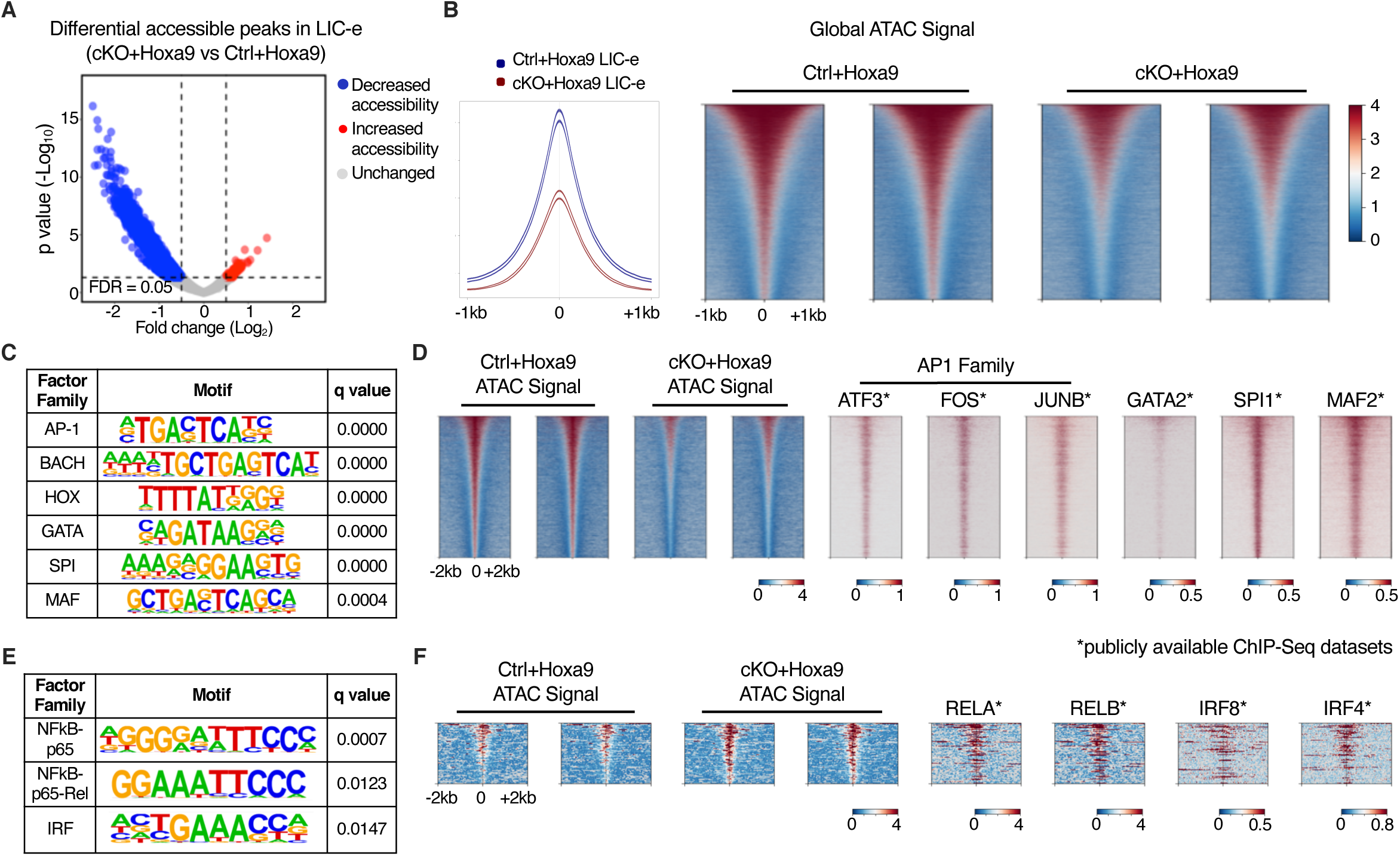
Effects of Phf6 loss on chromatin accessibility in LIC-e cells. **A,** Volcano plot showing differentially accessible regions in LIC-e cells from *cKO+Hoxa9 compared Ctrl+Hoxa9*. (n = 3 biological replicates) **B,** Representative signal profile (left) and metagene plots (right) showing genome-wide intensity of ATAC signal in *Ctrl+Hoxa9* and *cKO+Hoxa9* LIC-e cells. **C,** HOMER analysis for regions of decreased chromatin accessibility in *cKO+Hoxa9* LIC-e cells, showing enrichment of AP-1, HOX, GATA, SPI1, and MAF motifs. **D,** Representative metagene plots at regions of decreased chromatin accessibility in *cKO+Hoxa9* LIC-e, show ChIP-Seq signal for select proteins whose motifs are seen to be enriched through HOMER in (**C**). **E,** HOMER analysis for regions of increased chromatin accessibility in *cKO+Hoxa9* LIC-e cells showing enrichment of NF-kB and IRF motifs. **F,** Representative metagene plots at regions of increased chromatin accessibility in *cKO+Hoxa9* LIC-e, show ChIP-Seq signal for select proteins whose motifs are seen to be enriched through HOMER in (**E**). Publicly available ChIP-Seq datasets in leukemia or myeloid cells were used in metagene heatmaps (Table S3). All plots were centered around ATAC-Seq peaks. SeqPlots was used to draw all metagene plots.

## DISCUSSION

A majority of *PHF6* mutations in myeloid and lymphoid leukemias are frameshift and nonsense mutations scattered throughout the gene body, presumed to cause loss of gene function and indicating that PHF6 acts as a leukemia suppressor. In concordance with this genomic observation, multiple mouse studies have reported increased HSC self-renewal with *Phf6* knockout (17–19,27), and have reported enhanced T-ALL progression when *Phf6* knockout is combined with activating mutations in *Notch1* or *Jak3*, or overexpression of *Tlx3* (17,18,20). In contrast, a recent publication (Hou et al, (23)) reported that AML induced by *BCR-ABL*, *AML1-ETO*, and *MLL-AF9* fusions is impaired by *Phf6* loss, reaching the counterintuitive conclusion that *Phf6* is required for myeloid leukemogenesis. Our paper addresses this controversy, and we demonstrate that *Phf6* loss accelerates AML in a broadly-relevant model, and that it does so by increasing the frequency and persistence of leukemic stem cells, a finding that is harmonious with the known role of *Phf6* as a repressor of HSC self-renewal.

We first interrogated the BEAT AML dataset, which showed that *PHF6* mutations are associated with worsened survival in human AML. Then, by using the mouse *Hoxa9* transduction AML model, we demonstrated that *Phf6* knockout led to increased colony replating, increased disease burden *in vivo*, progressively worsened survival on serial transplantation, and increased LIC frequency. To our knowledge, AML hierarchy and LIC immunophenotypes had not been previously defined in the *Hoxa9* model, and we determined that a simplified gating scheme could identify an LIC-enriched population that we termed ‘LIC-e’, which was the only population capable of robust colony plating and engraftment. We found that *Phf6* loss led to an expansion of LIC-e cells, and that this expansion could be recapitulated *in vitro*, with *Phf6* knockout LIC-e cells showing the ability to persist in culture while wild-type LIC-e cells were depleted. Contrary to reports that *Phf6* loss alters cell cycle or apoptosis, we found evidence of neither, and instead found that *Phf6* loss leads LIC-e cells to produce more progeny with persistent LIC-e identity, thus indicating that *Phf6* specifically controls the balance between LIC self-renewal and differentiation. RNA-Seq and ATAC-Seq showed that *Phf6* loss skews the LIC-e transcriptome towards a more stem-like state, and leads to reduced accessibility at loci bound by AP-1, GATA2, and SPI1.

There could be multiple reasons for why our results stand in contrast to those of Hou et al. First, as stated earlier, we specifically picked *Hoxa9* transduction as a driver that broadly recapitulates AML biology, while Hou et al used fusions that are not known to co-occur with *PHF6* in patients. It is therefore unclear if their models reflected the *in vivo* context in which *PHF6* mutations gain a clonal advantage in humans. Second, though our group and theirs both used *Vav-Cre* to knock out *Phf6*, Hou et al used *Flox*-only mice as negative controls, while we used *Vav-Cre*-only mice. *Cre* toxicity has been reported with other *Cre* models (45,46), and may be a contributory factor to these discordant phenotypes; our approach eliminates *Cre* toxicity as a potential confounder. Third, it is highly likely that *Phf6* loss produces different downstream transcriptional effects in different epigenetic contexts; we ourselves observed that *Phf6* loss produces stemness-skewing changes only in LIC-e cells but not in committed cells, and it is therefore conceivable that *Phf6* loss produces divergent effects in certain AML contexts compared to others. Careful work in human genomics and mouse modeling will be required to define the co-mutational contexts in which *Phf6* loss is cooperative, neutral, or actively deleterious to AML stemness and growth.

We observed a striking reduction in global chromatin accessibility in LIC-e cells on *Phf6* loss, with a third of all peaks showing significantly reduced ATAC signal. Collectively, this led to reduced accessibility at sites with AP-1 family, GATA2, and SPI1 occupancy, and increased accessibility at a small number of sites with NF-kB and IRF family occupancy. This observation does not allow us to draw any immediate conclusion about the direct molecular action of PHF6 protein on chromatin, and we believe it to be a consequence of indirect effects that may cumulatively lead to reduced overall accessibility. We also acknowledge that the LIC-e population is not homogenous, and likely contains cells with higher or lower global chromatin accessibility. Future work using bulk and single-cell ATAC-Seq studies will be required to map out the precise chromatin effects of *Phf6* loss in the Hoxa9 model, as well as in other models such as co-mutation with *RUNX1* or *ASXL1*.

In summary, our work defines an LIC-enriched population in *Hoxa9*-driven AML, and the hierarchy through which it differentiates and expands to produce the bulk of the AML population. We show that *Phf6* loss increases the number of these LIC-e cells, not through increased cycling, but by producing progeny with persistent LIC-e identity. Taken together with the relatively pure phenotype of increased HSC self-renewal observed when *Phf6* is knocked out in homeostatic marrow, this presents a useful system to demonstrate how the normal process of self-renewal is co-opted in AML to drive the self-renewal of leukemic stem cells.

## MATERIALS AND METHODS

### Analysis of BEAT AML data

Mutational and survival information for 557 human adults with AML was obtained from the BEAT AML (26) dataset (through the cBioPortal website https://www.cbioportal.org/, RRID:SCR_014555). Patients were stratified based on ELN 2017 classification provided on cBioPortal. Kaplan-Meier survival curves were plotted separately for adverse and intermediate ELN risk groups (*PHF6* mutations were rare in favorable risk AML), and statistical significance was assessed using the Log-rank (Mantel-Cox) test.

### Mice

Cryopreserved sperm from *Phf6^fl/Y^* mice (serial# 4621-2 / G4621) was purchased from the Mouse Clinical Institute at GIE-CERBM (GIE-Centre Européen de Recherche en Biologie et Médecine, France), and pups were generated by the Children’s Hospital of Philadelphia Transgenic Core by IVF using C57BL/6J oocytes. *Vav-Cre* mice were originally generated by Thomas Graf (47) and were provided as a generous gift by Warren Pear (Department of Pathology, University of Pennsylvania Perelman School of Medicine). Both alleles were backcrossed with pure C57BL/6J for over 10 generations. All animals were maintained and experiments were carried out according to the University of Pennsylvania’s Animal Resources Center and IACUC protocols.

### *Hoxa9* overexpression construct and retrovirus preparation

pMSCV Hoxa9-IRES-GFP retroviral construct plasmid was a generous gift from Kathrin Bernt (Division of Oncology, Children’s Hospital of Philadelphia). Retroviruses were generated by transient co-transfection of 293T cells with retroviral vector and ecotropic packaging plasmids. Supernatant containing retroviruses was collected after 48 and 72 hrs of culture, filtered through a 0.45 μm filter, and used to transduce mouse bone marrow cells.

### *Hoxa9* transduction of mouse bone marrow

Age-matched *Vav-Cre^Cre/+^ Phf6^+/Y^* (*Ctrl*) or *Vav-Cre^Cre/+^ Phf6^fl/Y^* (*cKO*) littermate mice were intraperitoneally injected with 150 mg/kg 5-fluorouracil (5-FU, Fresenius Kabi, 101720), and bone marrow cells were isolated from femurs after 4 days. Red blood cells were lysed with ACK buffer (Lonza, #10-548E), and cells were cultured for 2 days in IMDM (Thermo Fisher Scientific, #12440053) with 15% fetal bovine serum (Gemini Bio, #100-100), 20 ng/ml mSCF (Peprotech, #250-03), 20 ng/ml mTPO (Peprotech, #315-14), 20 ng/ml mFLT3 ligand (Peprotech, #250-31L), 10 ng/ml mIL3 (Peprotech, #213-13), and 10 ng/ml mIL6 (Peprotech, #216-16). Cultured cells were then transduced with spinofection (2400 RPM, 90 min) with pMSCV Hoxa9-IRES-GFP retroviral supernatant and 4 ug/ml polybrene (Millipore Sigma, #TR-1003), and transferred to fresh media 18 hours after spin. GFP+ cells were purified after 2 days by fluorescence-activated cell sorting (BD FACSAria II).

### mRNA isolation and qPCR

To confirm that *Phf6* cKO does not express Phf6 mRNA, we isolated mRNA from bone marrow of 3 *Ctrl* and 3 *cKO* animals using TRizol (Thermo Scientific, #15596026) and prepared cDNA using 500 ng of total RNA using high capacity cDNA reverse transcriptase (Thermo Scientific, #4368813). qPCR for *Phf6* was carried out using primers TGTTTTCGTCTGCTTTGGTG (forward) and ATCATGCAATGCACAGTGGT (reverse), and for *Gapdh* (endogenous control) using primers CCCAGCTTAGGTTCATCAGG (forward) and GAATTTGCCGTGAGTGGAGT (reverse).

### Immunophenotypic analysis and cell sorting

Total bone marrow was isolated by flushing and crushing bones (femur, tibia, humerus, ilium and column vertebralis) in PBS supplemented with 2% FCS. Red blood cell lysis was performed for 5 min at room temperature in ACK lysing buffer (Lonza, #10-548E). For flow cytometry, a single-cell suspension of bone marrow cells was stained with the requisite panel of fluorochrome-conjugated antibodies on ice for 30 minutes to 1 hour. A complete list of antibodies is provided in Table S1. Cells were analyzed using an LSR Fortessa flow cytometer (BD Biosciences), and data were analyzed using FlowJo (FlowJo, LLC). When purification of cell populations was desired, cells were sorted using a BD Aria II sorter (BD Biosciences) into chilled PBS with 10% BSA for further processing.

### Colony Forming assay

For myeloid clonogenic progenitor assays, *Hoxa9*-transduced cells were plated on 35 mm Petri dishes (Corning) in a 1.1 mL culture mixture containing Methocult (STEMCELL Technologies, #M3234) with 20 ng/ul mIL3 (Peprotech, #213-13), 20 ng/ul mIL6 (Peprotech, #216-16), and 20 ng/ul mSCF (Peprotech, #250-03). Colonies propagated in culture were scored and replated (500 cells/plate) every 7 days. Average colony size was determined by counting the number of cells obtained per plate and dividing by the number of colonies. A total of 8 replating rounds were performed.

### *In vitro* culture of *Hoxa9*-transduced cells

*Hoxa9*-transduced bone marrow cells from *Ctrl* or *cKO* mice were sorted for GFP positivity, and cultured in IMDM (Thermo Scientific, #12440053) with 15% fetal bovine serum (Gemini Bio), 20 ng/ml mSCF (Peprotech, #250-03), 20 ng/ml mTPO (Peprotech, #315-14), 20 ng/ml mFLT3 ligand (Peprotech, #250-31L), 10 ng/ml mIL3 (Peprotech, #213-13), and 10 ng/ml mIL6 (Peprotech, #216-16). Cells were diluted to 100K/ml every other day.

### AML mouse model and serial bone marrow transplantation

For all transplantation experiments, recipient mice were wild-type C57BL/6J mice aged 8-20 weeks, lethally irradiated with 1100cGy administered as two split doses of 550cGy given 4 hours apart. *Hoxa9*-transduced bone marrow cells from *Ctrl* or *cKO* mice were injected retro-orbitally into primary recipient mice along with 400K radioprotectant cells of healthy age-matched bone marrow. Mice were monitored for activity and body conditioning score. For survival curve cohorts, mice were euthanized when humane endpoints for morbidity were met. For serial transplantation, bone marrow from primary recipient mice was harvested at 8 weeks after transplantation and lysed with ACK buffer. Sorted GFP+ cells along with 400K radioprotectant cells were retro-orbitally injected into lethally irradiated secondary recipients. Subsequent tertiary transplants were similarly performed using sorted GFP+ cells from marrow harvested from secondary recipients at 8 weeks after transplantation. Each cohort contained 8-20 week old age and sex-matched recipient mice.

### Limiting dilution transplantation assay

To calculate LIC (Leukemia initiating cell) frequency in freshly transduced marrow or in 8-week transplanted marrow, cohorts of lethally irradiated recipient mice were transplanted with 400K, 100K, 30K, 3K or 1K GFP+ cells from the respective source. Mice were euthanized when humane endpoints for morbidity were met or when >375 days had passed (experiment termination), and analyzed for the presence of GFP+ cells in bone marrow. All mice that achieved humane endpoints for morbidity had >80% GFP+ cells in marrow, and were counted as responders to the corresponding dose. At experiment termination (>375 days), a small number of mice with >1% GFP were also counted as responders (most had nearly 80% GFP positivity), while mice with <1% GFP+ cells in marrow and peripheral blood were counted as non-responders. LIC frequency was calculated using ELDA software (48).

### Histopathology

Mouse femur, spleen, liver, and lungs were harvested and fixed in formaldehyde. Femurs were decalcified and tissue paraffin blocks and slides were prepared and stained by the University of Pennsylvania Molecular Pathology and Imaging Core. Bone marrow was cytospun and peripheral blood was smeared onto glass slides, following which the slides were air-dried and stained using HEMA 3 stain (Thermo Fisher Scientific, #22-122911). Slides were imaged using a Leica microscope.

### *In vivo* cell cycle analysis

The BD Pharmingen™ APC BrDU Kit (BD Biosciences, #552598) was used as per the manufacturer’s protocol. 1 mg/mouse BrdU (BD bioscience, #552598) was administered intraperitoneally at 8 weeks after primary transplant, and bone marrow was harvested after 2 hours. Following ACK lysis, cells were stained for surface antigens, fixed and treated with DNase to expose BrdU antigen, and incubated with Pacific Blue conjugated anti-BrdU antibody as per the instruction manual. Flow cytometry was performed as described above.

### *In vitro* cell cycle analysis

For cell cycle analysis and to calculate the length of the cell cycle, the Click-iT™ EdU Pacific Blue™ Flow Cytometry Assay kit (Invitrogen™, #C10418) was used as per the manufacturer’s protocol. Briefly, 50K sorted LIC-e cells were incubated for 10 hours in media containing 10 µM EdU. Cells were harvested every hour, washed with 1% BSA in PBS, and stained for surface antigens. Cells were then fixed using the Click-iT fixative for 15 minutes at room temperature, and subsequent washes were performed using the 1X Click-iT saponin-based permeabilization and wash reagent. Cells were incubated for 30 minutes at room temperature in the dark with the Click-iT reaction cocktail composed of CuSO_4_, Pacific Blue dye, reaction buffer additive and PBS, following the manufacturer’s instructions. Flow cytometry was performed as described above.

For EdU pulse-chase experiments, 50K purified LIC-e cells were incubated for 2 hours in media containing 10 µM EdU (pulse). Cells were washed 3 times with fresh media to remove EdU, and were cultured for an additional 8 hours (chase) before being harvested and stained as described above.

### Apoptosis

Apoptotic and necrotic cell analysis was performed using Pacific Blue™ Annexin V Apoptosis Detection Kit with 7-AAD (BioLegend, #640926) as per manufacturer’s instructions. Cells were distinguished as live (AnnexinV-, 7AAD-), apoptotic (AnnexinV+, 7AAD-), necrotic (7AAD+).

### RNA sequencing

50K LIC-e and 150K committed leukemic cells were sorted from bone marrow of primary recipients of *Ctrl+Hoxa9* and *cKO+Hoxa9* cells at 8 weeks after transplantation. Total RNA was extracted using TRIzol LS reagent (Thermo Fisher Scientific, #10296010) and RNeasy Mini Kit (Qiagen, #74106). Total RNA concentration and RNA integrity number was determined using a Bioanalyzer (Agilent Technologies) and high sensitivity RNA Screentape (Agilent, #5067-5579). Samples with RNA integrity number 8 or higher were used for RNA library preparation. For LIC-e cells, RNA libraries were prepared using Smarter Stranded Total RNA-Seq Kit - Pico Input Mammalian (Takara Bio, #635005). For committed cells, RNA libraries were prepared using NEBNext Ultra RNA Library Prep Kit for Illumina (New England Biolabs, #E7530L). All libraries were prepared using manufacturer’s instructions. Libraries were quantified using high sensitivity D1000 Screentape (Agilent Technologies, #5067-5584) and KAPA Library Quantification Kit (Thermo Fisher Scientific, #KK4824). Equimolar libraries from each sample were pooled and subjected to 75-bp paired-end sequencing on a NextSeq 550 machine (Illumina). Raw reads were demultiplexed and FASTQ files were generated using Bcl2fastq v2.20.0.422 (Illumina). The reads were trimmed for low quality/adapter sequence using Trimmomatic (RRID:SCR_011848) (49), followed by aligning to the mouse genome (mm10) with STAR v2.7.10b aligner (50). Gene-level read counts were generated using the featureCounts tool from Rsubread v2.0.3) (51). Read count normalization and differential gene expression testing was performed using DESeq2 v1.30.1 (52). GSEA analysis was performed using DESeq2 normalized read counts using GSEAv4.1.0. Mac application (SeqGSEA, RRID:SCR_005724, Broad institute). Gene signatures were downloaded from the MSigDB (Broad) or from the GSE datasets directly in the GSEA application. Gene set enrichment data was represented using default parameters and an FDR cutoff of 0.05.

### ATAC sequencing

After 4 days of *Hoxa9* transduction, 50K LIC-e cells were sorted from *Ctrl+Hoxa9* and *cKO+Hoxa9* bone marrow cultures. Cells were lysed and their nuclei were isolated, followed by tagmentation performed using Nextera Tn5 transposase (Illumina Nextera ATAC seq kit, 20034197). Libraries were prepared using dual-end indexing using NEBNext Ultra II DNA Library Prep Kit (New England Biolabs, #E7645L) using the manufacturer’s protocol. The size and concentration of each sample’s library were determined using Agilent 2200 Tapestation and KAPA library quantification kit (Roche, #KK4824) respectively. Libraries from all samples were pooled in equimolar concentrations and denatured along with 1% of PhiX for paired-end sequencing. The pooled library was quantified using KAPA library quantification kit (Roche KK4824) and equimolar libraries were pooled and subjected to 75-bp paired-end sequencing on a NextSeq 550 machine (Illumina). Raw reads were demultiplexed and FASTQ files were generated. Reads were trimmed of contaminating adapter sequences using Trimmomatic (49) and aligned to the mouse genome (mm10) using Bowtie2 (RRID:SCR_016368) (53). Peaks of transposase-accessible chromatin were called and quantified using MACS2 v2.2.7.1 (54). Differential and consensus peak analyses were performed using the DiffBind v3.0.15 package in R software (RRID:SCR_012918). Motif analysis was also performed for all regions with significant differential accessibility after filtering out low-confidence regions using HOMER (55). Public ChIP-Seq datasets (Table S3) for ATF3, FOS, JUN, GATA2, SPI1, MAF2, IRF4, IRF8, RELA and RELB were mapped to mm10 genome and used for generating metagene plots using SeqPlot software.

### Statistical Analysis

Statistical analysis was performed using GraphPad Prism v9.5.1 (GraphPad Software). We assumed equal distribution of variance between groups and performed either 1-way Anova with Sidak’s multiple comparison testing or Student’s *t-test*. Kaplan-Meier curves were used to represent survival, where significance was calculated with the log-rank (Mantel-Cox) test. We considered results p < 0.05 as statistically significant and represented using *p < 0.05, **p < 0.01, ***p < 0.001; ****p < 0.0001. We analyzed serial limited dilution leukemia-initiating cell data using the ELDA tool(48).

## Blinding

All histopathological analyses were performed in a blinded fashion.

## Replication

All transplant experiments were performed in multiple cohorts of recipients, and data were compiled for final analyses.

## Power Analysis

Power analyses for mouse experiments were at 90% confidence, significance level of 0.05.

## Inclusion and Exclusion Criteria

N/A, no clinical trial data in this study

## Attrition

N/A, no clinical trial data in this study

## Cell Line Authentication

N/A, no cell lines used in this study

## DATA AVAILABILITY

All generated datasets have been deposited to GEO: GSE249108 for RNA-Seq on LIC-e cells, GSE249109 for RNA-Seq on committed leukemic cells, and GSE249110 for ATAC-Seq datasets on LIC-e cells.

## ACKNOWLEDGEMENTS

We thank Nancy Speck, Ivan Maillard, Wei Tong, and Kathrin Bernt for helpful discussions. V.R.P. is supported by National Institute of Health (NIH) grant R01-HL155144 (NHLBI), American Cancer Society (ACS) grant 129784-IRG-16-188-38-IRG, an American Society of Hematology (ASH) Faculty Scholar Award, and a University of Pennsylvania Covid-19 Research Disruption Mitigation Fund. S.S.J. is supported by an ASH Restart Award. C.A is supported by a Scholar Award from the American Society of Hematology (ASH) and Co-Operative Center for Excellence in Hematology (CCEH) grant by the National Institute of Diabetes and Digestive and Kidney diseases (NIDDK). Data for this manuscript were generated in the Penn Cytomics and Cell Sorting Shared Resource Laboratory at the University of Pennsylvania and is partially supported by the Abramson Cancer Center NCI Grant (P30 016520). The research identifier number is RRid: SCR_022376.

## AUTHOR CONTRIBUTIONS

V.R.P. conceived the project and supervised the study. S.S.J. designed and performed all experiments. S.S.J., A.P. and V.R.P. wrote the manuscript with input from all authors. S.S.G., S.S.J. and A.P performed bioinformatic analyses, including scripting for downstream analysis and graphical representation. S.S.J., C.A., and A.P. made and edited figures. S.G. performed histological characterization and provided histopathology images. J.G. contributed to breeding and maintaining the mouse colony.

---

**Figure S1.**
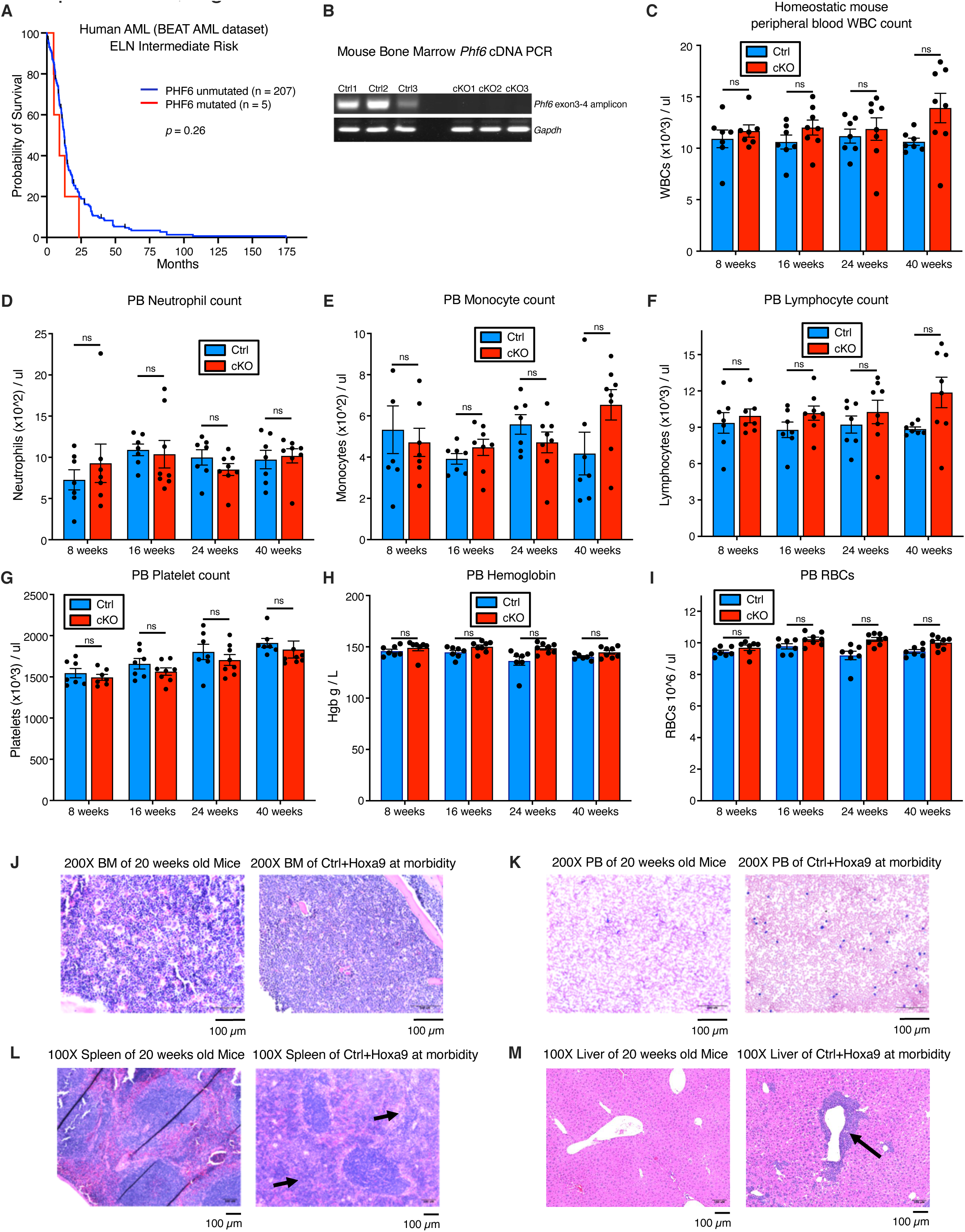
(related to Figure 1). **A,** Kaplan-Meier survival curves for *PHF6* mutated and unmutated intermediate risk (ELN classification) adult AML patients from the BEAT AML dataset. Statistical significance was calculated using the Log-rank (Mantel-Cox) test. **B,** *Phf6* mRNA expression in bone marrow of *Vav-Cre^Cre/+^ Phf6^+/y^* (*Ctrl*) and *Vav-Cre^Cre/+^ Phf6^fl/y^* (*cKO*) mice. *Gapdh* is shown as a loading control. n = 3 biological replicates **C-I,** Bar graphs depicting peripheral blood counts for WBCs, neutrophils, lymphocytes, monocytes, platelets, hemoglobin and RBCs for *Ctrl* and *cKO* mice at 8, 16, 24, and 40 weeks of age. n = 8 biological replicates **J,** H&E staining of bone marrow collected from a representative *Ctrl+Hoxa9* mouse at morbidity, with age-matched homeostatic (non-leukemic) WT mouse shown for reference. Scale bar is 100um at 200X. **K,** Wright-Giemsa stain of blood smears from a representative *Ctrl+Hoxa9* mouse at morbidity, with age-matched homeostatic WT mouse shown for reference. Scale bar is 100um at 200X. **L-M,** H&E staining of (**L**) spleen and (**M**) liver collected from a representative *Ctrl+Hoxa9* mouse at morbidity, with age-matched homeostatic (non-leukemic) WT mouse shown for reference. Scale bar is 100um at 100X. Arrows indicate leukemic infiltration. All bar graphs show mean ± SEM. n.s. = non-significant by student t-test.

**Figure S2.**
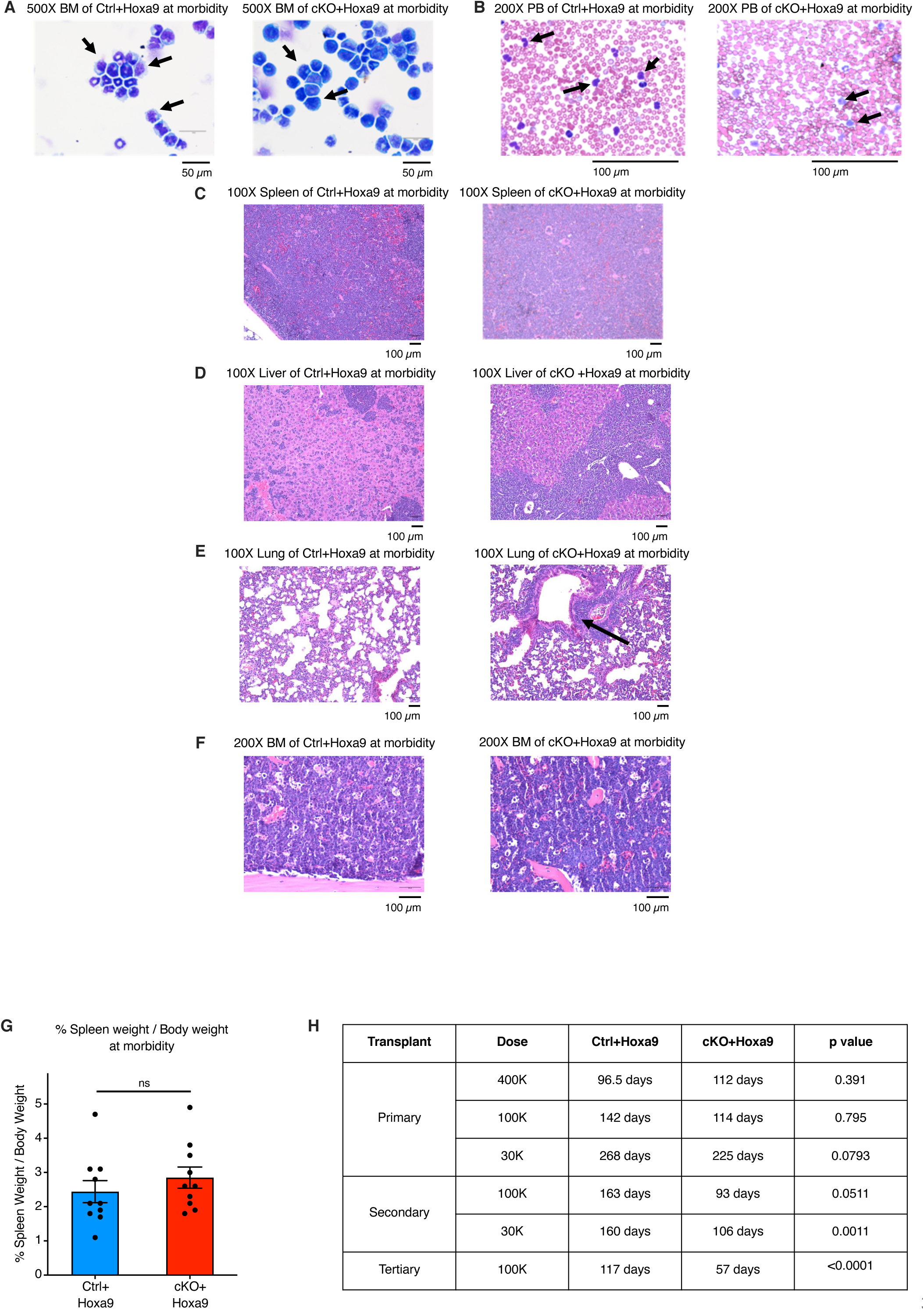
(related to Figure 1). Loss of Phf6 accelerates HoxA9-driven AML on serial transplantation **A,** Representative image of Wright-Giemsa stain of bone marrow cytospins of *Ctrl+Hoxa9* and *cKO+Hoxa9* mice at morbidity. Scale bar is 50um at 500X. Arrows indicate blast cells. **B,** Representative image of Wright-Giemsa stain of blood smears of *Ctrl+Hoxa9* mouse and *cKO+Hoxa9* mouse at morbidity. Scale bar is 100um at 200X. Arrows indicate blast cells. **C-F,** Representative image of H&E staining of (**C**) spleen, (**D**) liver, (**E**) lung, and (**F**) bone marrow collected from *Ctrl+Hoxa9* mouse and *cKO+Hoxa9* mouse at morbidity. Scale bar is 100uM at 100X. Black arrows indicate leukemic infiltration. **G,** Bar graph showing percent spleen / body weight at morbidity. Statistical significance was calculated using the Student t test. n.s. = non-significant by student t-test. **H,** Table with median survival and statistical significance for primary, secondary and tertiary transplantation with different dose of *Ctrl+Hoxa9* and *cKO+Hoxa9* cells.

**Figure S3.**
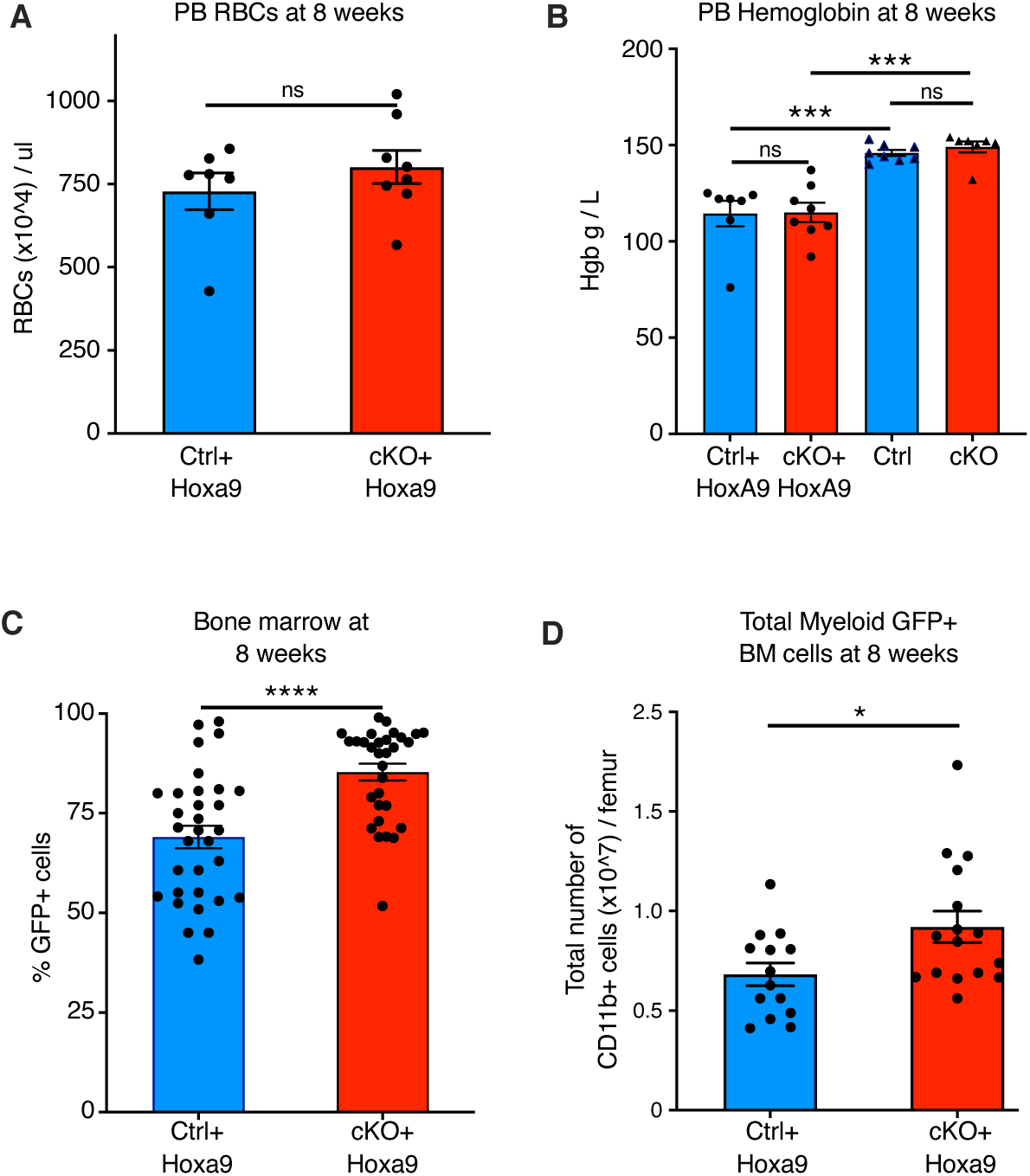
(related to Figure 2). *Phf6* loss increases leukemic disease burden **A-B,** Bar graphs of peripheral blood counts for (**A**) RBCs and (**B**) hemoglobin level of *Ctrl+Hoxa9* and *cKO+Hoxa9* mice at 8 weeks, post primary transplantation. Hemoglobin levels of homeostatic (non-leukemic, non-transplanted) *Ctrl* and *cKO* mice are shown for reference. n = 7 biological replicates, n.s. = non-significant by student t-test. **C-D,** Frequency of (**C**) GFP+ cells per femur and (**D**) total number of myeloid GFP+ cells per femur in the bone marrow of primary recipient after 8 weeks of transplantation of *Ctrl+Hoxa9* and *cKO+Hoxa9* cells.

**Figure S4.**
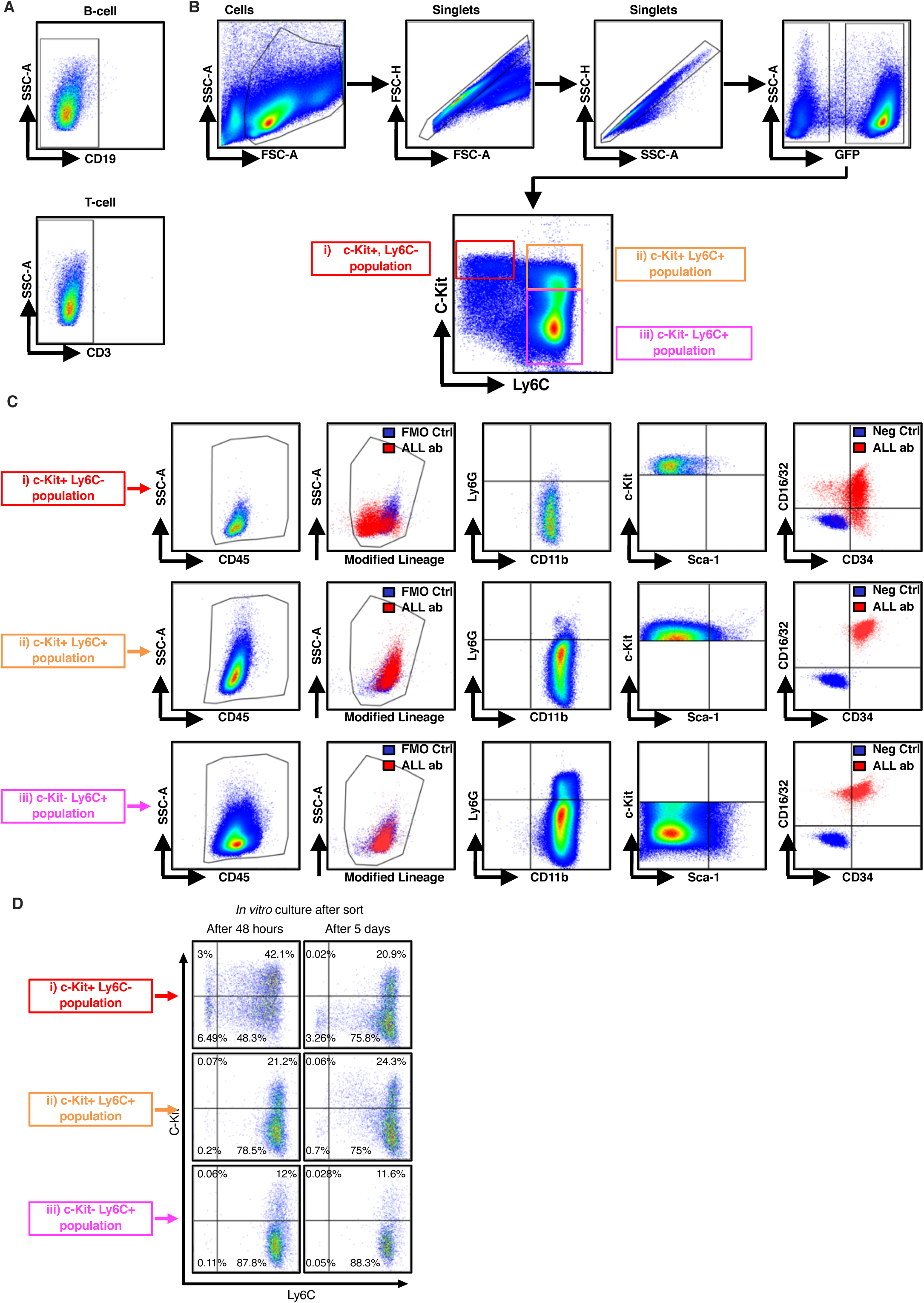
(related to Figure 3). Phenotypic characterization of *Hoxa9*-driven AML subpopulations **A,** Representative FACS plot of bone marrow leukemic cells (GFP+) from primary recipient mice at 8 weeks after transplantation. FACS plots demonstrate absence of B cell specific (CD19) and T cell specific (CD3) antigen expression. **B-C,** FACS plots demonstrating the gating strategy used to compartmentalized GFP+ cells. We have designated the three sub-populations: red (c-Kit+ Ly6C-), orange (c-Kit+ Ly6C+) and pink (c-Kit- Ly6C+) as LIC-e, Committed and Differentiated leukemic cells respectively. Each population is further characterized for the expression of CD45 (pan hematopoietic marker), modified non-myeloid lineage cocktail (CD3, CD19, CD45R, Ter119, CD49b), CD11b (myeloid marker), Ly6G (neutrophil marker), c-kit, Sca-1, CD34, and CD16/32. LIC-e cells were: mLin^-^Kit^+^Sca^-^ CD11b^dim^ Ly6G^-^ CD34^+^ CD16/32^-/+^ Ly6C^-^ Committed cells were: mLin^-^Kit^+^Sca^-^ CD11b^+^ Ly6G^-/+^ CD34^+^ CD16/32^+^ Ly6C^+^ Differentiated cells were: mLin^-^Kit^-^Sca^-^ CD11b^+^ Ly6G^-/+^ CD34^+^ CD16/32^+^ Ly6C^+^ Note: The main flow plot in Fig S4B is also shown in main Fig 3A (shown here with detailed gating strategy). **D,** Representative flow cytometry plots showing immunophenotypes of cells obtained after 48 hours and 5 days of *in vitro* culture of the three sorted subpopulations of *Ctrl+Hoxa9* transplanted marrow shown in Fig 3A.

**Figure S5.**
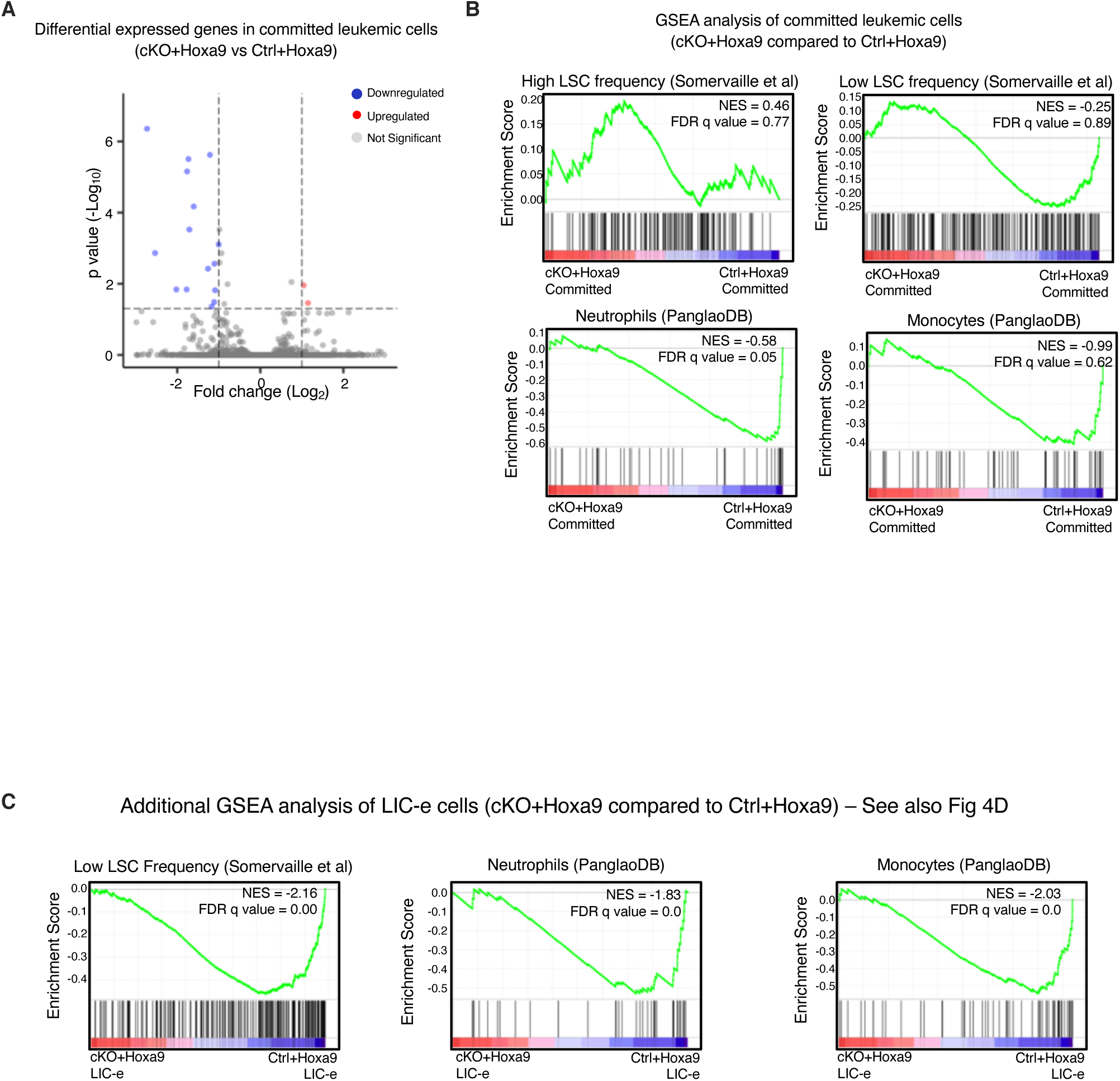
(related to Figure 4). *Phf6* loss has minimal effects on the transcriptome of the committed (c-Kit+, Ly6C+) AML cell population **A,** Volcano plot showing differentially expressed genes between sorted committed cells (c-Kit+, Ly6C+) from *Ctrl+Hoxa9* and *cKO+Hoxa9* mice at 8 weeks after transplantation. Minimal gene expression changes were seen, with 28 downregulated and 3 upregulated genes in *cKO+Hoxa9* compared to *Ctrl+Hoxa9*. **B,** Gene set enrichment analysis (GSEA) of the *cKO+Hoxa9* committed cell transcriptome compared to *Ctrl+Hoxa9* for gene sets related to high and low LSC frequency, mature neutrophils, and mature monocytes. Normalized Enrichment scores (NES) and FDR q values are shown. **C,** Gene set enrichment analysis (GSEA) of the *cKO+Hoxa9* LIC-e transcriptome compared to *Ctrl+Hoxa9* for gene sets related to low LSC frequency, mature neutrophils, and mature monocytes. Normalized Enrichment scores (NES) and FDR q values are shown.

**Figure S6.**
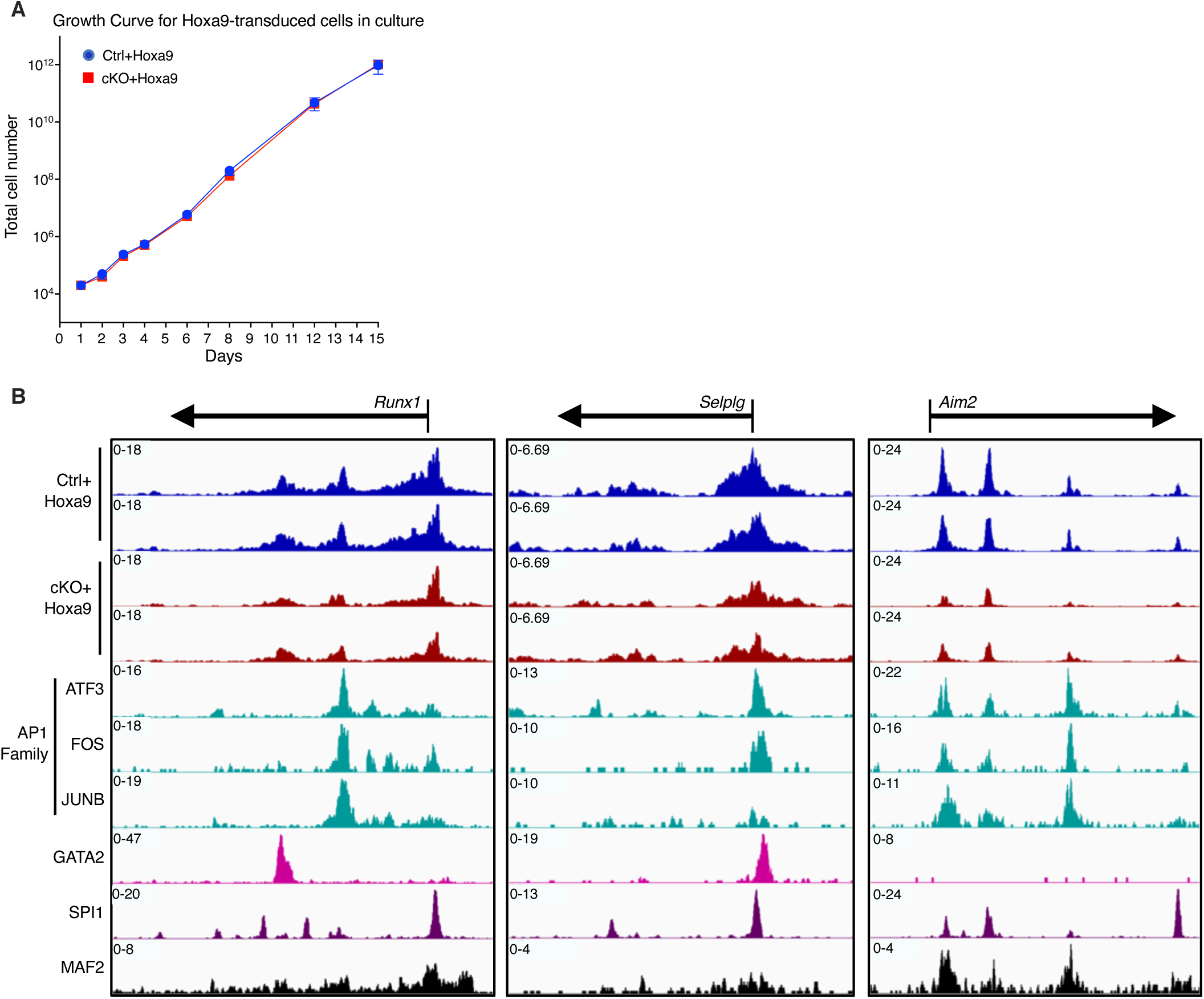
(related to Figure 6). Tracks of transcription factor occupancy at differentially expressed genes **A,** Growth curve of *in vitro* cultured *Ctrl+Hoxa9* and *cKO+Hoxa9* cells. **B,** IGV browser tracks of ATAC-Seq signal in LIC-e cells at promoters of genes downregulated or unchanged in *cKO+Hoxa9* along with ChIP-Seq signal of proteins whose motifs are enriched in regions of reduced accessibility in *cKO+Hoxa9* LIC-e *cells*. *Ctrl+Hoxa9* and *cKO+Hoxa9* peaks are scaled to the same Y-axis. Transcription factor ChIP-Seq tracks are taken from publicly available datasets in myeloid or leukemic cell types (see Table S3).

